# Evaluating sequencing strategies for endometrial microbiome profiling in endometrial cancer: a comparative study of short- and long-read 16S rRNA approaches

**DOI:** 10.1101/2025.09.02.673865

**Authors:** Sophia Bebelman, Anastasiia Artuyants, Bianca Nijmeijer, Sandra Fitzgerald, Claire Henry, Cherie Blenkiron

## Abstract

**Background:** Endometrial cancer (EC) is the most common gynaecological malignancy globally, with rising incidence and notable disparities in outcomes. In New Zealand, EC rates have increased significantly, particularly among Māori and Pacific women, who face higher risks of advanced disease and poorer outcomes. Microbial dysbiosis has been implicated in EC pathogenesis, but characterising the uterine microbiome is challenging due to low microbial biomass and high contamination risk.

**Aims:** This study aimed to pilot a protocol that could inform the preparation of a larger cohort trial. Short-read Illumina MiSeq and long-read Oxford Nanopore Technologies (ONT) 16S rRNA gene sequencing were investigated to profile the uterine microbiome in people with EC.

**Methods and Results:** Uterine and vaginal swabs were analysed to assess platform performance in terms of DNA recovery, sequencing success, diversity metrics, and taxonomic resolution. The impact of sample freezing or immediate lysis prior to DNA extraction was also evaluated. ONT sequencing provided enhanced species-level resolution and improved detection of low-abundance taxa but showed variable performance in low-yield samples. Freezing prior to cell DNA extraction modestly increased bacterial 16S copy numbers and improved community consistency. Contamination was a problem across both platforms, particularly in low-biomass samples, but can be minimised during data analysis.

**Conclusion:** This study provides practical guidance for sequencing platform selection and sample handling in uterine microbiome research. Our findings support future efforts to elucidate microbial contributions to EC pathogenesis and highlight the importance of rigorous contamination control. Importantly, this is the first presentation of a New Zealand cohort and contributes valuable data from an underrepresented population and informs future research in diverse clinical settings.

## 1. Introduction

Endometrial cancer (EC) is the most common gynaecological malignancy in high-income countries, with incidence rates rising globally.^1^ It accounts for approximately 95% of uterine cancers and is typically diagnosed in postmenopausal women.^2^ While early-stage EC is often treatable, advanced or recurrent disease presents a significant clinical challenge. Strikingly, in New Zealand (Aotearoa), EC incidence has increased by nearly 60% over the past decade, with Māori and Pacific women disproportionately affected.^3^ These populations experience higher rates of aggressive disease, delayed diagnosis, and poorer outcomes, highlighting the need for improved understanding of EC pathogenesis and more inclusive and global research efforts.

Advances in high-throughput sequencing have transformed our understanding of the human microbiome and its role in immunity, metabolism, and disease.^4^ In oncology, microbial dysbiosis has been linked to colorectal, cervical, and breast cancers, with growing interest in its potential contribution to EC pathogenesis.^5, 6, 7^

The uterus, once considered a sterile environment, is now recognised to host a unique and dynamic microbial community, albeit in relatively lower abundance than adjacent reproductive tract regions. Emerging evidence suggests that microbial imbalance may influence local inflammation, immune modulation, and carcinogenesis.^8, 9, 10, 11^ However, research into the uterine microbiome remains limited, largely due to technical challenges.^10, 12^ A key challenge lies in the low microbial biomass of endometrial samples, which complicates accurate taxonomic classification and increases susceptibility to contamination from reagents, handling, or adjacent anatomical sites.^13^

16S rRNA gene sequencing is widely used as an affordable approach for bacterial community profiling, but platform selection influences resolution and reliability.^14^ ^15^ Short-read Illumina MiSeq sequencing offers high accuracy and robust bioinformatics support, but is limited by its short read length, which restricts genus- and species-level classification.^16, 17, 18^ ^19^ In contrast, long-read Oxford Nanopore Technologies (ONT) sequencing enables full-length 16S profiling, potentially improving taxonomic resolution and detection of low-abundance taxa.^16, 19^ An advantage of Nanopore technology is its potential to generate sequence data from low-input samples, making it suitable for studies with limited biological material.^20^

Despite these technological advances, few studies have systematically compared these sequencing platforms for low-biomass uterine samples, and none have focused on EC patients in New Zealand - a geographically isolated population with distinct demographic and clinical characteristics. Moreover, methodological variables such as sample preservation and DNA extraction protocols may further influence microbial community profiles, yet these factors are rarely evaluated in low-biomass environments.

In this study, we conducted a direct comparison of Illumina MiSeq and ONT 16S rRNA sequencing on uterine swabs from EC patients. Our objectives were: (1) to evaluate platform performance in terms of DNA recovery, sequencing success, diversity metrics, and taxonomic resolution; and (2) to assess the feasibility of uterine sampling and identify methodological considerations for future microbiome research in EC. By addressing both technical and population-specific considerations, this work will inform future uterine microbiome research and support efforts to elucidate microbial contributions to EC pathogenesis.

## 2. Methods

### 2.1 Ethical approval and patient recruitment

A total of 31 participants were recruited from Te Whatu Ora Wellington Hospital for the collection of uterine swabs and genetic material analysis. Ethics approval was obtained from the New Zealand Health and Disability Ethics Committee (HDEC 15/CEN143) and the University of Otago (H20/002 and H25/0421). Eligible participants were aged 18 years or older and undergoing hysterectomy for any stage of endometrial cancer. Exclusion criteria included pregnancy, breastfeeding, and recent antibiotic use.

In a separate cohort, vaginal swabs were collected from three individuals undergoing surgery for suspected endometriosis at Te Whatu Ora Wellington Hospital. Ethics approval for this study was granted by the Central Health and Disability Ethics Committee (HDEC 2022 EXP 12616). Participants were aged between 16 and 45 years, with exclusions for menstruation at the time of sampling and known cervical abnormalities.

### 2.2 Sample collection and processing

Following surgical removal of the uterus, four swabs were collected per participant - two from the tumour site and two from adjacent non-tumorous tissue. In cases where the tumour size limited visible delineation of the tissues, fewer swabs were collected.

To identify the optimal sample collection method for downstream analysis, four swab and storage protocols were evaluated, varying in swab type, buffer composition, and storage temperature: Method A: Copan FLOQ (Copan Diagnostics) swabs stored at room temperature (RT) in DNA/RNA Shield lysis buffer (Zymo Research) for up to 4 weeks; Method B: Zymo DNA/RNA Shield collection swabs stored at RT; Method C: FLOQ swabs collected in Tris-EDTA (TE) buffer (Thermo Fisher Scientific), centrifuged, and the pellet resuspended in DNA/RNA Shield (RT); Method D: FLOQ swabs stored in TE buffer at −80°C (for 1-2 months), thawed, centrifuged, and the pellet resuspended in lysis buffer prior to processing.

Vaginal swabs were collected by the operating surgeon or nurse following anaesthesia and prior to surgical preparation. The collection medium consisted of 1x phosphate-buffered saline (PBS; Thermo Fisher Scientific) supplemented with 25 mM trehalose dihydrate (Sigma-Aldrich) and 25 mM HEPES (Thermo Fisher Scientific). Swabs were obtained by inserting a FLOQ swab into the vagina and rotating for approximately 30 seconds, then placed into a tube containing 1 mL of collection medium. Samples were visually inspected, and any showing signs of blood contamination were excluded.

All swabs were preserved in their respective buffers and transported to the University of Auckland for subsequent analysis.

### 2.3 DNA extraction and quantification

Genomic DNA was extracted using a ZymoBIOMICS™ DNA Microprep Kit (Zymo Research), with minor modifications to enhance microbial lysis. Briefly, 1 mL of swab/cell lysate (in DNA/RNA shield) was transferred to BashingBead™ Lysis Tubes and subjected to mechanical disruption using a Disruptor Genie bead beater (Scientific Industries) at maximum speed for 20 minutes. Lysates were centrifuged at 10,000 x g for 1 minute, and 400 µL of supernatant was mixed with DNA Binding Buffer and applied to Zymo-Spin™ IC Columns. Columns were washed with DNA Wash Buffers 1 and 2, and DNA was eluted in 20 µL of DNase/RNase-Free Water. To remove potential inhibitors, eluates were purified using the Zymo-Spin™ II-μHRC Filter and ZymoBIOMICS™ HRC Prep Solution by centrifugation at 16,000 x g for 3 minutes, and stored at −20°C until analysis. DNA concentrations were measured using the Qubit™ dsDNA High Sensitivity (HS) Assay Kit (Thermo Fisher Scientific), following the manufacturer’s protocol and measured using the Qubit 4 Fluorometer.

### 2.4 Droplet digital PCR (ddPCR)

To assess the ratio of human to bacterial DNA, droplet digital PCR (ddPCR) was performed using the QX200™ Droplet Digital PCR System (Bio-Rad). Dual assays targeting the human *NRAS* gene and the bacterial 16S rRNA gene were used.

Each 23 µL ddPCR reaction contained 1x ddPCR Supermix for Probes (no dUTP) (Bio-Rad), 0.43 µM each of *NRAS* forward (5′-TGGTGAAACCTGTTTGTTGG-3′) and reverse (5′-ATTGGTCTCTCATGGCACTG-3′) primers, 0.43 µM each of universal bacterial forward (5′-TGGAGCATGTGGTTTAATTCGA-3′) and reverse (5′-TGCGGGACTTAACCCAACA-3′) primers, 0.21 µM of a FAM-labelled 16S probe (5′ 6-FAM–CACGAGCTGACGACARCCATGCA–ZEN–Iowa Black FQ 3′), 0.21 µM of a HEX-labelled NRAS probe (5′ HEX–CTGGATACA–ZEN–GCTGGACAAGAAG–IABkFQ 3′), and 5 ng of DNA template (0.22 ng/µL final concentration).

Droplets were generated using a Bio-Rad Automated Droplet Generator and then amplified using a Bio-Rad C1000 Touch Thermal Cycler (Bio-Rad) under the following conditions: enzyme activation at 95 °C for 10 minutes; 40 cycles of 94 °C for 30 seconds and 59 °C for 1 minute; followed by enzyme deactivation at 98 °C for 10 minutes and read using a QX200 Droplet Reader. Absolute quantification was performed using QuantaSoft™ Analysis Software (v1.7.4.0917). Positive controls included *NRAS*-only (human cells) and *E. coli*-only samples; a no-template control (NTC) was also included.

### 2.5 16S rRNA gene amplicon sequencing (Illumina MiSeq)

DNA samples were submitted to Auckland Genomics for 16S rRNA gene amplicon sequencing. DNA concentrations were normalised to 5 ng/µL prior to library preparation. The ZymoBIOMICS™ Microbial Community DNA Standard (Zymo Research) was used as a positive control, and molecular-grade water served as a negative control.

Amplicon libraries targeting the V3-V5 hypervariable regions were prepared using the Illumina protocol with modifications. The following primers were used: 357F: 5′– TCGTCGGCAGCGTCAGATGTGTATAAGAGACAGCCTACGGGAGGCAGCAG–3′ and 926R: 5′– GTCTCGTGGGCTCGGAGATGTGTATAAGAGACAGCCGTCAATTCMTTTRAGT–3′.

PCR amplification was performed using Platinum™ SuperFi II PCR Master Mix (Thermo Fisher Scientific) for 35 cycles, using 2.5 μL (12.5 ng) of DNA with the following thermal cycling conditions: an initial denaturation step at 95 °C for 3 minutes, followed by 35 cycles of denaturation at 95 °C for 30 seconds, annealing at 55 °C for 30 seconds, and extension at 72 °C for 30 seconds. A final extension was carried out at 72 °C for 5 minutes. Amplicons were purified using AMPure XP magnetic beads and eluted in 20 µL of nuclease-free water.

A second indexing PCR was performed using custom Nextera XT index primers, followed by another bead cleanup and elution in 40 µL. Indexed libraries were pooled equimolarly, assessed for fragment size using a Bioanalyzer, normalised to 4 nM, spiked with ∼24% PhiX, and sequenced on an Illumina MiSeq using a 600-cycle v3 kit (2×300 bp).

A total of 14,744,455 read pairs passed the filter, generating approximately 7.58 Gbp of data, with an average of ∼150,000 paired-end reads per sample, providing sufficient depth for microbial community analysis.

### 2.6 Full-length 16S rRNA gene sequencing (Oxford Nanopore Technologies)

Genomic DNA samples were submitted to Auckland Genomics, University of Auckland, for full-length 16S rRNA gene sequencing using Oxford Nanopore Technologies (ONT). DNA concentrations were normalised to 2.5 ng/μL, and 10 μL (25 ng) was used for the PCR.

First-Round PCR Amplification: Full-length 16S rRNA gene regions were amplified using ONT-adapted universal primers targeting positions 27F to 1492R, with overhang sequences compatible with downstream barcoding: ONT_27F: 5′– TTTCTGTTGGTGCTGATATTGCAGAGTTTGATCCTGGCTCAG–3′ and ONT_1492R: 5′–ACTTGCCTGTCGCTCTATCTTCTACGGYTACCTTGTTACGACTT–3′.

Amplification was performed using Platinum™ SuperFi II PCR Master Mix (Thermo Fisher Scientific,) with the following thermal cycling conditions: an initial denaturation step at 95 °C for 1 minute, followed by 35 cycles of denaturation at 95 °C for 20 seconds, annealing at 55 °C for 30 seconds, and extension at 65 °C for 2 minutes. A final extension was carried out at 65 °C for 5 minutes.

Positive controls included the ZymoBIOMICS™ microbial community DNA standard (Zymo Research), and molecular-grade nuclease-free water served as the negative control. Amplicons were purified using a 1:1 ratio of AMPure XP beads and eluted in 20 µL of nuclease-free water. Post-cleanup, amplicons were quantified and re-normalised to 2.5 ng/μL.

Indexing PCR and Library Preparation: Barcoding was performed using LongAmp® Hot Start Taq 2X Master Mix (New England Biolabs) and a EXP-PBC096 barcoding kit from ONT. Reactions were set up in 25 µL total volume, with the following thermal cycling conditions: an initial denaturation at 95 °C for 3 minutes, followed by 12 cycles of denaturation at 95 °C for 15 seconds, annealing at 62 °C for 15 seconds, and extension at 65 °C for 45 seconds. A final extension step was performed at 65 °C for 5 minutes.

Barcoded samples were pooled at equimolar concentrations, purified using AMPure XP beads, and eluted in 40 µL of nuclease-free water. Libraries were prepared using an SQK-LSK114 ligation sequencing kit (ONT), following the manufacturer’s instructions.

Sequencing and Basecalling: Sequencing was performed on a PromethION FLO-PRO114M flow cell. During sequencing, barcode trimming was disabled, and adapter trimming was enabled. Basecalling was conducted in two stages using Dorado (v0.8.0): Real-time basecalling during the run using the “high accuracy” model and Post-run basecalling using the “super high accuracy” model to improve read quality.

### 2.7 Data analysis

#### 2.7.1 Illumina MiSeq Data Processing

Illumina reads were processed using the QIIME2 (2024.2 amplicon distribution) pipeline via the New Zealand eScience Infrastructure.^21^ Only forward reads were used (covering 16S rRNA section V3), trimmed to 200 bp, and denoised using DADA2.^22^ A phylogenetic tree was constructed using the align-to-tree-mafft-fasttree workflow from the q2-phylogeny plugin.

Taxonomic classification was performed using a pre-trained naïve Bayes classifier via the q2-feature-classifier plugin.^23^ The classifier was trained on full-length 16S sequences from the SILVA 138 OTU database at 99% similarity.^24, 25, 26, 27^ For comparison, taxonomy was also assigned using the NCBI database, ^28^ with a classifier generated using RESCRIPt, filtered for bacterial sequences with a minimum read length of 800 bp.^25^

#### 2.7.2 Oxford Nanopore Technologies (ONT) Data Processing

ONT long-read data were processed using the Epi2me 16S workflow, also run via the New Zealand eScience Infrastructure. Taxonomic classification was performed to the genus level using minimap2 and the SILVA 138.1 database, with default settings unless otherwise specified.

Minor differences in taxonomic group naming between SILVA database versions used for Illumina and ONT were corrected to match the most recent version prior to comparative analysis.

#### 2.7.3 Diversity analysis in R

Both Illumina and ONT datasets were further analysed using R (version 4.3.2).^29^ For alpha rarefaction, genus-level data were converted into phyloseq objects, and rarefaction curves were generated using the rarecurve function from the vegan package (v2.6-10).^30^

For diversity analysis, datasets were converted into phyloseq objects and rarefied to a sequencing depth determined by alpha rarefaction analysis (Figures S2 and S9).^18, 31^ Alpha diversity metrics (observed richness, Shannon, and Simpson indices) were calculated using estimate_richness from phyloseq, and results were aggregated using dplyr (v1.1.4).^32^ Wilcoxon tests were used for pairwise comparisons, and Kruskal-Wallis tests followed by Dunn’s tests with Benjamini-Hochberg correction were applied for multi-group comparisons using rstatix (v0.7.2).^33^ Visualisations were generated using ggplot2 (v3.5.2).^34^

Beta diversity was assessed using Bray-Curtis dissimilarity and principal coordinate analysis (PCoA) via phyloseq (v1.46.0). PERMANOVA testing was performed using adonis2 from vegan, and pairwise PERMANOVA was conducted using pairwise.adonis2 from the pairwiseAdonis package (v0.4.1), with false discovery rate (FDR) controlled via the Benjamini–Hochberg method.^35^

#### 2.7.4 Taxonomic composition and PCA in R

For taxonomic bar plots, genus-level data were filtered to exclude unassigned reads. Where genus-level classification was unavailable, higher taxonomic ranks were used. The top 29 genera were selected based on read counts, and the remaining taxa were grouped under “Other”. Relative abundance was calculated per sample and visualised using ggplot2.

For principal component analysis (PCA), genus-level data were Hellinger-transformed using decostand from vegan, and taxa with zero variance were removed. Dimensionality reduction was performed using prcomp, and the percentage of variance explained by each principal component was calculated. The first two components were visualised using ggplot2.

## 3. Results

### 3.1 DNA yield and sequencing success

To evaluate sequencing platform performance in profiling the uterine microbiome, endometrial tumour swabs were collected from 26 patients undergoing hysterectomy for endometrial cancer (Figure 1a and 1b). This enabled direct sampling of the uterine cavity, minimising contamination from the lower genital tract. From each patient, two swabs were taken from the tumour site and two from adjacent non-cancerous endometrial tissue. In total, 112 samples were obtained, representing an initial cohort that included 3 Māori, 2 Pacific peoples, and the remainder of European (Pākehā) descent. Comparative analysis revealed no significant differences in read count, microbial diversity, or taxonomic composition between tumour and adjacent tissue swabs (data not shown) in line with a prior study.^10^ Therefore, 1-2 swabs per patient from the tumour site only were retained for platform comparison to reduce redundancy.

**Figure 1.**
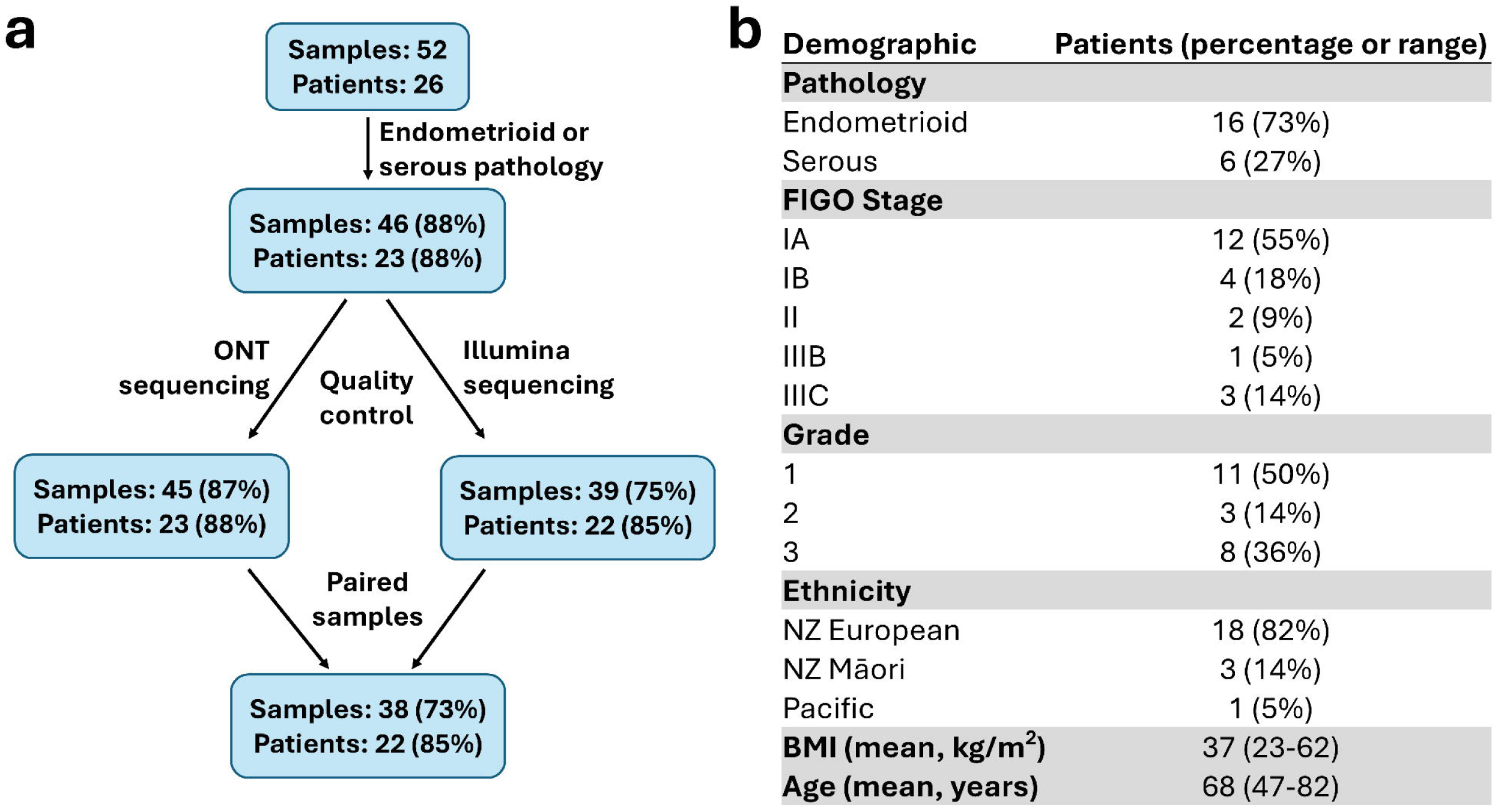
Sample and patient retention through processing and quality control. (a) Flow diagram showing the number and percentage of samples and patients retained after each stage of sequencing data processing and quality control. (b) Summary of demographic characteristics of patients whose samples passed all processing and quality control steps. Ethnicity is self-reported. Pathology and demographics were taken from clinical notes. Age and BMI are at surgery.

Three patients were excluded due to tumour histology inconsistent with the study’s focus on serous and endometrioid subtypes. Among the remaining 23 patients, 73% (n=17) presented with endometrioid carcinoma, consistent with its higher prevalence in the general population (Figure 1b). This subtype is frequently associated with metabolic comorbidities and tends to occur in younger women.^36^ This was reflected in the cohort’s average BMI of 37 kg/m² and the presence of five women under 60 years of age (Figure 1b).

To confirm the presence of bacterial DNA and assess sample suitability for sequencing, droplet digital PCR (ddPCR) was performed prior to library preparation (Table S1). The average human-to-bacterial DNA ratio was 166:1, consistent with the low microbial biomass expected in uterine samples. For comparison, high biomass vaginal swabs had an average human-to-bacteria ratio of 1:5.5. Despite the dominance of host DNA in uterine swabs, bacterial DNA was detectable in all samples.

A total of 46 swabs were sequenced using both ONT and Illumina platforms and distinct data analysis pipelines. Samples were excluded if they failed to meet quality thresholds defined by Epi2me (ONT) or QIIME2 (Illumina), including a minimum of 10,000 high-quality reads. ONT sequencing yielded higher read counts (mean = 677,000 reads) compared to Illumina (mean = 110,000 reads). The success rate for Illumina (75%) was lower than for ONT (87%). Only samples that passed QC for both platforms were included in downstream comparative analyses to ensure consistency, leaving 38 samples from 22 patients.

### 3.2 Taxonomic resolution and read depth

To assess taxonomic assignment performance between sequencing platforms, we evaluated two widely used reference databases: SILVA and NCBI. SILVA is a highly curated database optimised for accurate classification at higher taxonomic levels (e.g., phylum to genus), but it is not recommended for species-level classification when using Epi2me due to limited resolution. In contrast, the NCBI database supports species-level assignments in both Epi2me and QIIME2 pipelines, offering finer taxonomic granularity.^21, 25^

Initial analyses were conducted using both databases to compare taxonomic assignment, diversity metrics, and overall microbial profiling (Figures S1, S3, and S7). While NCBI enabled species-level resolution, SILVA demonstrated superior consistency and accuracy at higher taxonomic ranks and outperformed NCBI when benchmarked against a mock microbial community. Given the known limitations of 16S rRNA gene sequencing for species-level identification,^19^ and the greater reliability of SILVA at the genus level, all downstream comparative analyses between ONT and Illumina platforms were conducted using the SILVA database, with taxonomic resolution restricted to the genus level.

Taxonomic assignment efficiency was quantified across hierarchical levels for both sequencing platforms (Figure 2). ONT sequencing assigned approximately 80% of reads consistently from phylum to genus level, reflecting the advantage of long-read sequencing by spanning multiple variable regions of the 16S gene. Illumina sequencing, which targets a shorter hypervariable region, showed higher assignment rates at broader taxonomic levels - 95% of reads were classified from phylum to order. However, assignment rates declined at finer levels, with around 90% of reads classified at the family level and only 70% at the genus level. This drop-off in resolution is consistent with the limitations of short-read sequencing, which may not capture sufficient sequence variation for confident genus-level classification.

**Figure 2.**
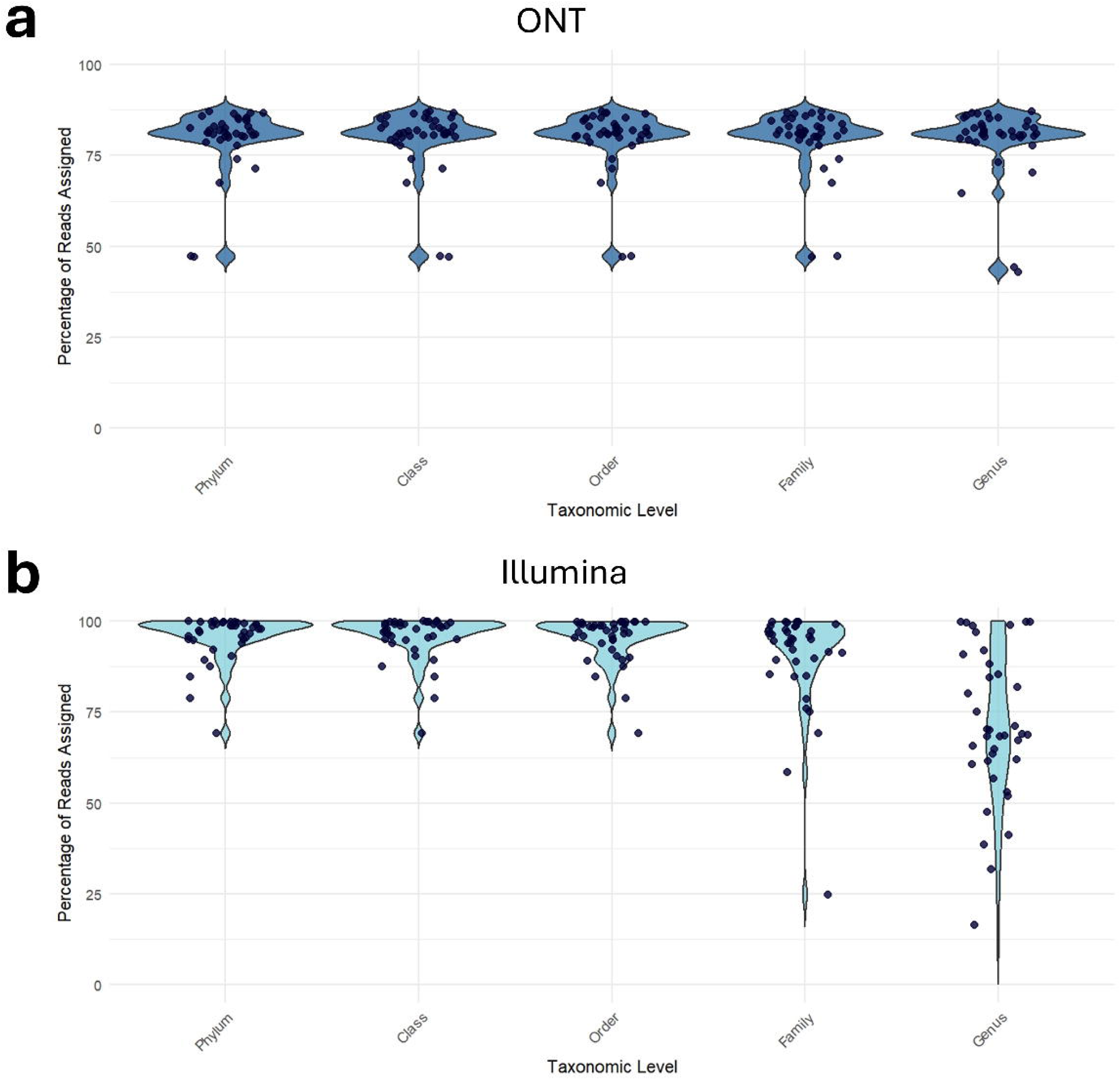
Taxonomic assignment efficiency between sequencing platforms. Violin plots showing the percentage of reads assigned at each taxonomic level for samples sequenced using (a) ONT and (b) Illumina platforms. Each dot represents an individual swab sample. Reads classified as unknown, unassigned, eukaryotic, or uncultured were excluded from the assigned read count to focus on confidently classified microbial taxa.

### 3.3 Alpha and beta diversity: genus richness, evenness, and community composition

To ensure comparability across sequencing platforms, alpha rarefaction was performed to normalise read depth and reduce bias in diversity estimates. Due to substantial differences in sequencing output, ONT samples yielded significantly higher read counts than Illumina, platform-specific rarefaction thresholds were applied: 140,000 reads for ONT and 10,000 reads for Illumina, balancing retention of microbial diversity with quality control (Figure S2).

Alpha diversity metrics revealed platform-specific differences in microbial richness and evenness. ONT samples exhibited significantly higher genus richness, with a mean of 53.3 observed genera per sample, compared to 18.4 genera in Illumina samples (Wilcoxon test, p = 4.21E-13; Figure 3a). This suggests that ONT’s long-read sequencing enables broader detection of microbial taxa.

**Figure 3.**
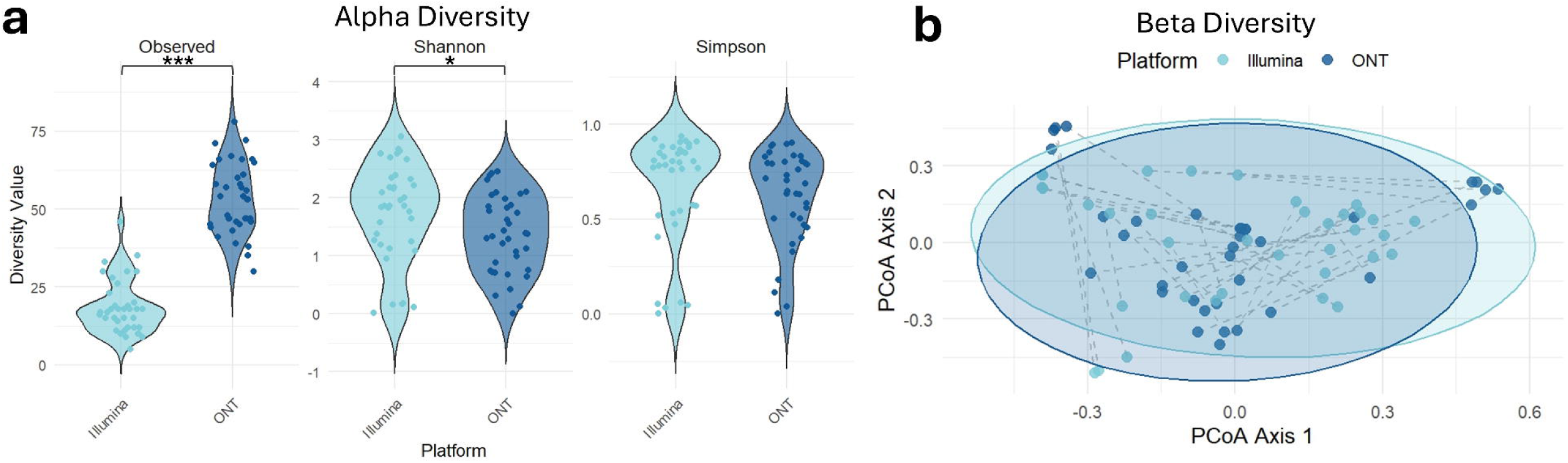
Comparison of alpha and beta diversity between Illumina and ONT sequencing platforms at the genus level. (a) Alpha Diversity Comparison: Violin plots of Observed, Shannon, and Simpson indices at the genus level, comparing Illumina (light blue) and ONT (dark blue) sequencing platforms. Each dot represents an individual sample. Samples were rarefied to 10,000 reads (Illumina) or 140,000 reads (ONT), excluding unassigned reads. Statistical significance was assessed using the Wilcoxon rank-sum test (*p*LJ<LJ0.05, \*\**p*LJ<LJ0.001). (b) Beta Diversity (Bray-Curtis PCoA): Principal coordinate analysis of Bray-Curtis dissimilarity at the genus level, comparing Illumina and ONT platforms. Each dot represents a sample; dotted lines connect matched samples sequenced on both platforms. Ellipses indicate 95% confidence intervals. Samples were rarefied as above, excluding unassigned reads. PERMANOVA yielded FLJ=LJ10.273, *p*LJ=LJ0.001 (999 permutations).

Interestingly, Shannon diversity indices, which account for both richness and evenness, were slightly higher in Illumina samples (mean = 1.74) than ONT (mean = 1.37, p = 2.46E-2). This indicates that although ONT detected more genera, Illumina reads were more evenly distributed across taxa. Simpson indices, which emphasise dominant taxa and community evenness, were similar between platforms (mean = 0.673 for Illumina, 0.619 for ONT, p = 7.98E-2), suggesting comparable evenness among the most abundant genera.

Beta diversity analysis was conducted to assess differences in microbial community composition between platforms. Bray-Curtis dissimilarity revealed substantial overlap in community profiles, yet the sequencing platform accounted for 12.8% of the observed variance, which was statistically significant (p = 0.001; Figure 3b). While many paired samples clustered closely, platform-specific effects were evident in some cases, highlighting the influence of sequencing technology on community structure.

### 3.4 Community structure differences and taxonomic resolution

Overall, there was strong concordance in taxonomic profiles between samples sequenced using Illumina and ONT platforms (Figure 4, S6). Microbial community composition was more influenced by inter-patient variability than by sequencing platform, with replicate swabs from the same patient showing high similarity regardless of the technology used. This suggests that both platforms reliably capture the dominant microbial signatures present in individual samples.

**Figure 4.**
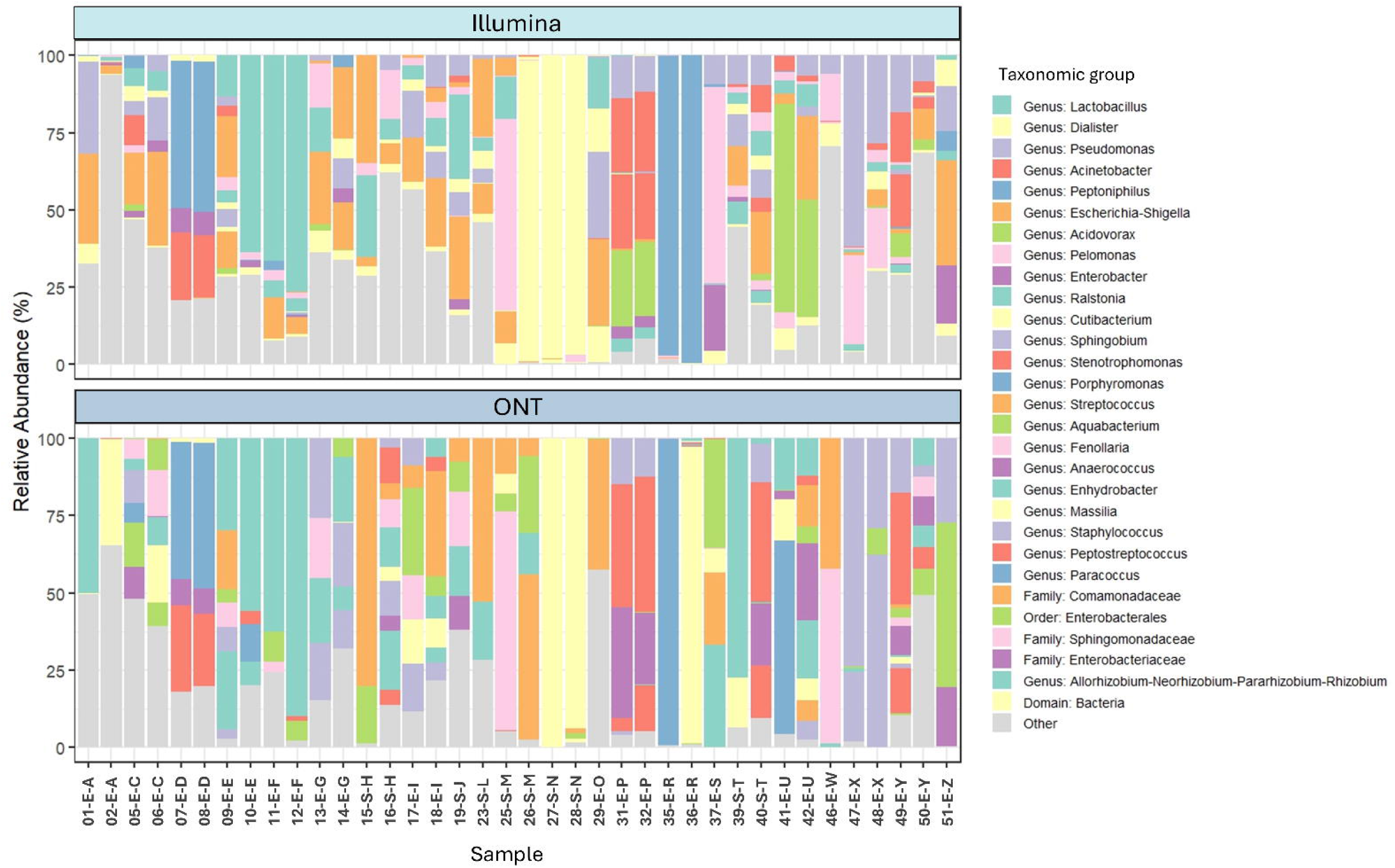
Relative abundance of taxonomic groups across samples. Bar plots showing the relative abundance of the 29 most abundant taxonomic groups across all samples. Taxonomic classification is presented at the genus level, or at a higher level when genus-level assignment was not possible. Unassigned reads were excluded from the analysis, and less abundant groups were combined into a single category labelled “Other”. Samples are represented by sample number, histological subtype (E - endometrioid; S - serous) and patient letter.

Despite overall concordance, differences in taxonomic resolution between platforms were evident. As previously noted (Figure 2), ONT assigned a greater proportion of reads at the genus level compared to Illumina. This discrepancy was particularly pronounced in taxa where Illumina reads were frequently classified only at the family or order level due to limitations in short-read resolution.

An example of this is the *Comamonadaceae* family, where ONT sequencing identified 23 distinct genera, while Illumina detected only 4. Notably, 79% of *Comamonadaceae* reads in Illumina samples were assigned only at the family level, with no genus-level classification. In contrast, ONT assigned just 0.003% of reads to the family level, with the majority classified into specific genera: *Acidovorax* (40%), *Pelomonas* (36%), and *Aquabacterium* (20%). *Acidovorax* was not detected at all by Illumina, and *Pelomonas* and *Aquabacterium* were underrepresented (16% and 3%, respectively).

These findings highlight the ability of ONT methodology to provide a more granular view of microbial community structure, particularly in low-biomass environments where taxonomic resolution is critical. The enhanced genus-level classification offered by ONT may be especially valuable for identifying subtle microbial shifts or rare taxa that could be biologically relevant but are missed or unclassified by short-read platforms. This has important implications for future studies aiming to link microbial signatures with clinical outcomes or disease mechanisms in endometrial cancer.

### 3.5 Detection of potential contaminants

To benchmark diversity metrics and validate sequencing performance, high-biomass vaginal swabs were included as positive controls. This group included 1 Māori and 2 of European (Pākehā) descent. The vaginal microbiome is typically dominated by *Lactobacillus* spp., which play a key role in maintaining mucosal health and preventing infection.^37^ Compared to uterine samples, vaginal swabs exhibited lower genus richness, lower Shannon and Simpson indices, and greater compositional similarity across samples (Figure S4a). Beta diversity analysis confirmed that vaginal samples formed a distinct cluster, clearly separated from uterine samples, with PCA further supporting this distinction (Figures S4b and S5).

To assess potential contamination, negative controls were included at two critical stages of the workflow: during DNA extraction and processing (processing control) and during library preparation and sequencing (no-template control). The Illumina negative controls yielded insufficient reads for meaningful analysis and were excluded from downstream comparisons.

In contrast, both ONT negative controls generated substantial read counts (506,090 and 250,240 reads, compared with 14,689–1,249,523 reads for ONT uterine swabs), which permitted detailed taxonomic profiling. Principal component analysis indicated that the no-template control clustered with several uterine samples (Figure S8). Taxonomic profiles of the controls showed distinct signatures: *Enhydrobacter*, *Cutibacterium* and *Acinetobacter*, taxa typically linked to skin or environmental sources, were identified in the sequencing control, while the processing control was dominated by *Lactobacillus*, consistent with possible carryover from vaginal samples processed in parallel. Because these taxa were also observed in some uterine samples, the control profiles were used to guide interpretation of low-abundance signals and to ensure confidence in distinguishing sample-specific from background contributions.

These findings highlight the need to distinguish true biological signals from background contamination in environments with low microbial load. With the implementation of appropriate controls, statistical tools can aid in contaminant identification and removal during analysis.^38, 39^ Despite challenges with contamination, ONT sequencing demonstrated clear advantages for microbiome profiling in the low-biomass uterine swabs, providing greater sequencing depth, broader taxonomic coverage, and a higher proportion of reads classified at the genus level, enabling a more detailed and nuanced view of the microbial communities present in the uterine environment.

### 3.6 Sample collection and storage method optimisation

We next evaluated sample collection and storage methods to optimise DNA yield and microbial recovery. The swabs used for platform comparison were collected using Method B (Figure 5), which involved immediate lysis of swab contents and storage at room temperature to simplify shipping. While this method was sufficient for sequencing, it consistently yielded low DNA concentrations, with low bacterial proportions, limiting downstream analysis.

**Figure 5.**
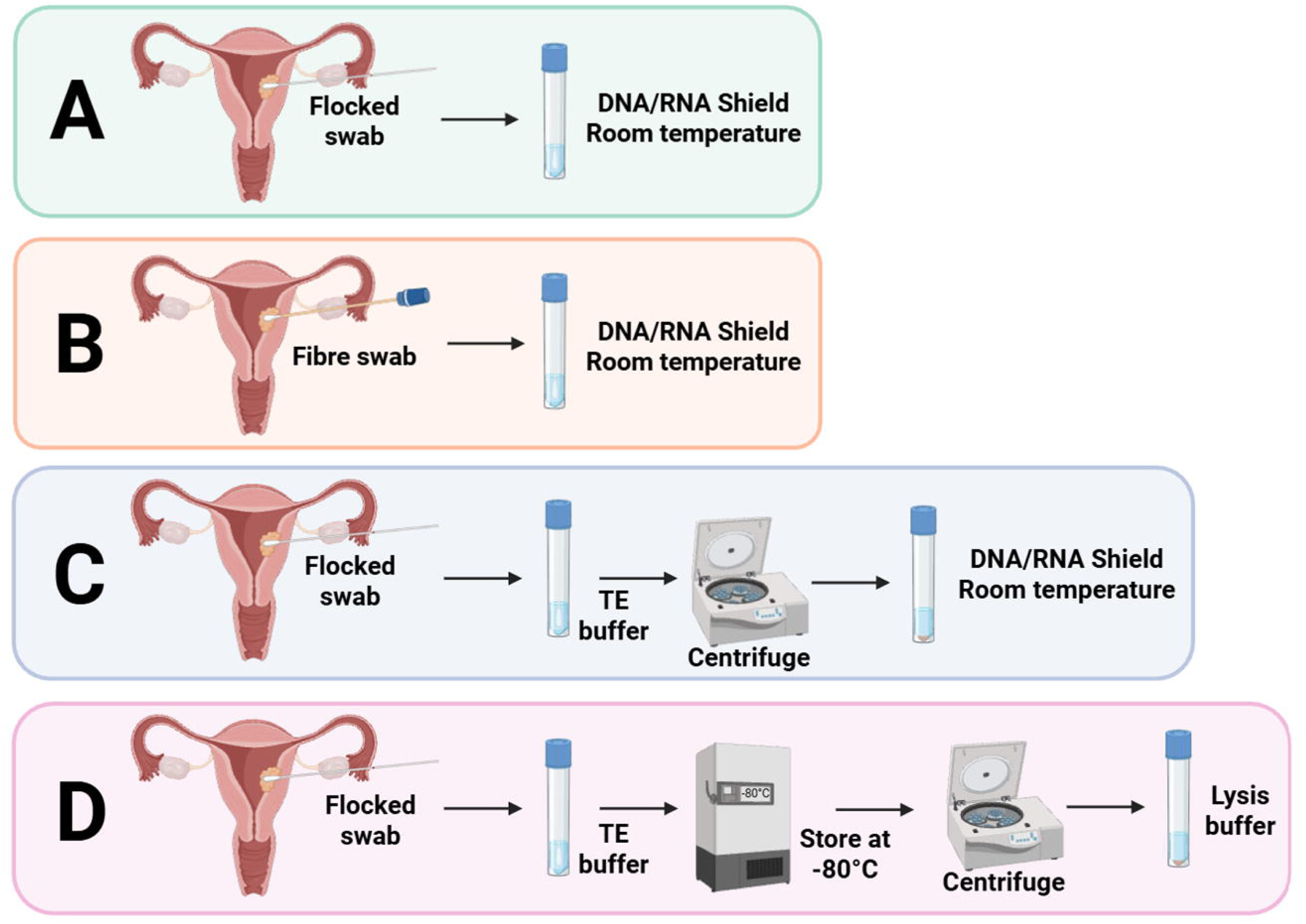
Overview of sample processing methods for uterine swab microbiome analysis. Illustration summarising the four sample processing methods (A-D) compared in this study for uterine swab microbiome analysis. Each method varies in its approach to sample handling, DNA extraction, and preparation. Figure created using BioRender.

To improve sample quality, we tested flocked swabs, which are used clinically and designed to enhance sample uptake and release, particularly in low-biomass environments. In addition to swab type, we assessed two alternative processing approaches. First, we lysed a cell pellet in DNA/RNA shield with ambient storage immediately after collection, which was designed to concentrate microbial DNA and reduce host DNA contamination (Method C). We also assessed freezing the swab immediately after collection in TE buffer, followed by thawing and pellet isolation prior to DNA extraction (Method D). This method was informed by a study suggesting that freezing can preserve microbial integrity and improve DNA recovery in low-biomass samples.^40^

Four endometrial tumour swabs from each of five patients were processed using Methods A–D and evaluated using the same quality control criteria applied to the initial cohort. DNA yield and human-to-bacteria ratios, as quantified by ddPCR, were consistent across all methods (data not shown). However, bacterial DNA copy numbers varied between methods and patients (Figure 6). Methods A and B showed PCR inhibition at the highest 50ng DNA input. Overall, Method D had a higher bacterial copy number and no evidence of PCR inhibition at higher input quantities.

**Figure 6.**
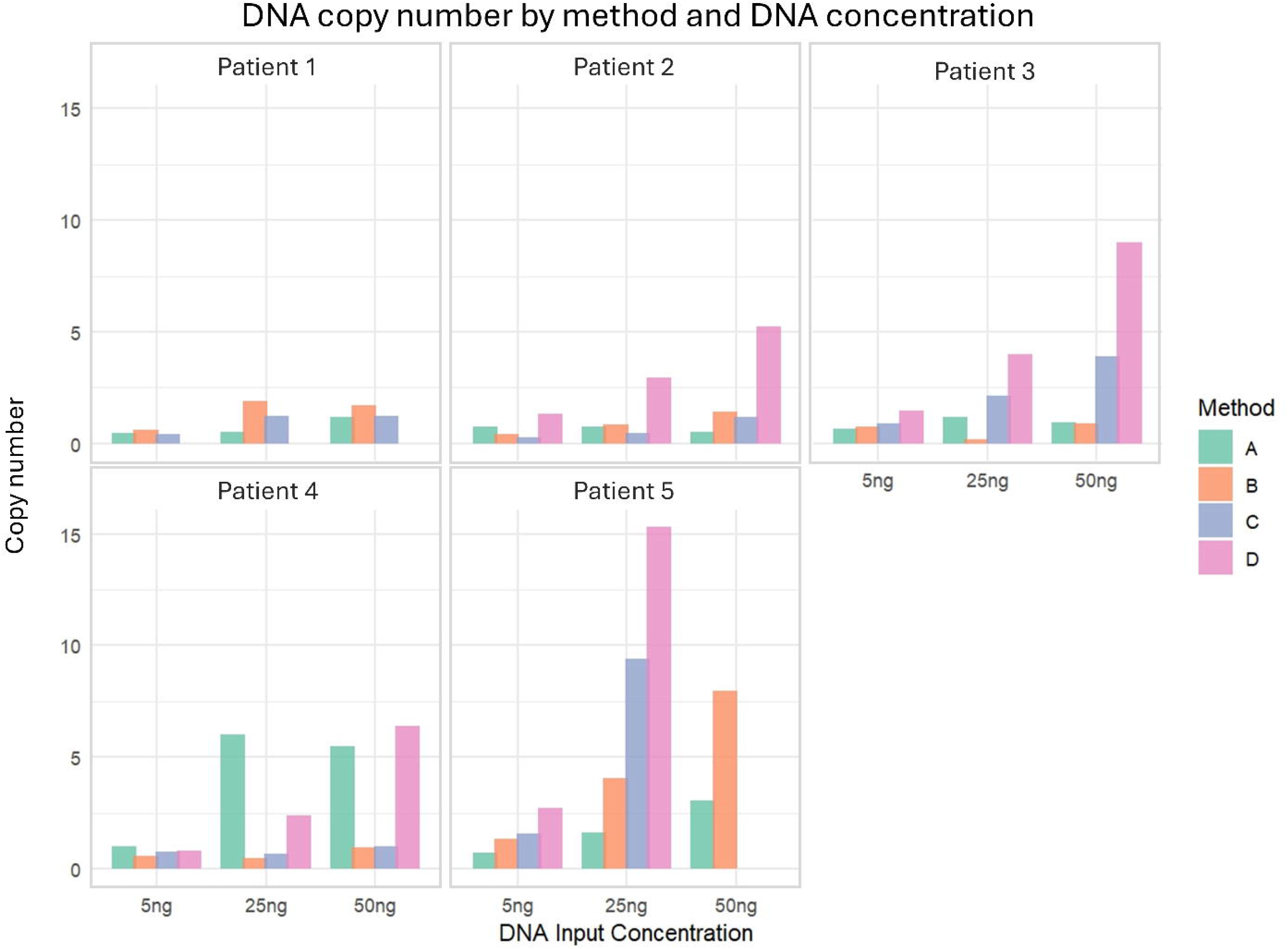
Bacterial 16S rRNA gene copy number across patients and processing methods. Bar plots showing bacterial 16S rRNA gene copy number across samples from patients 1-5, each collected using four processing methods (A-D). For each sample, three DNA input concentrations (5 ng, 25 ng, and 50 ng) were tested using ddPCR. Each bar represents an individual sample. Missing bars are due to insufficient sample for ddPCR to be carried out.

To assess the impact of these methods on microbial diversity, all samples were rarefied to 130,000 reads to normalise sequencing depth (Figure S9). Alpha diversity analysis revealed no statistically significant differences between methods in terms of observed genus richness, Shannon, or Simpson indices (Figure 7a). However, Methods C and D showed greater variability in the number of genera identified, suggesting that these approaches may be more sensitive to patient-specific microbial profiles.

**Figure 7.**
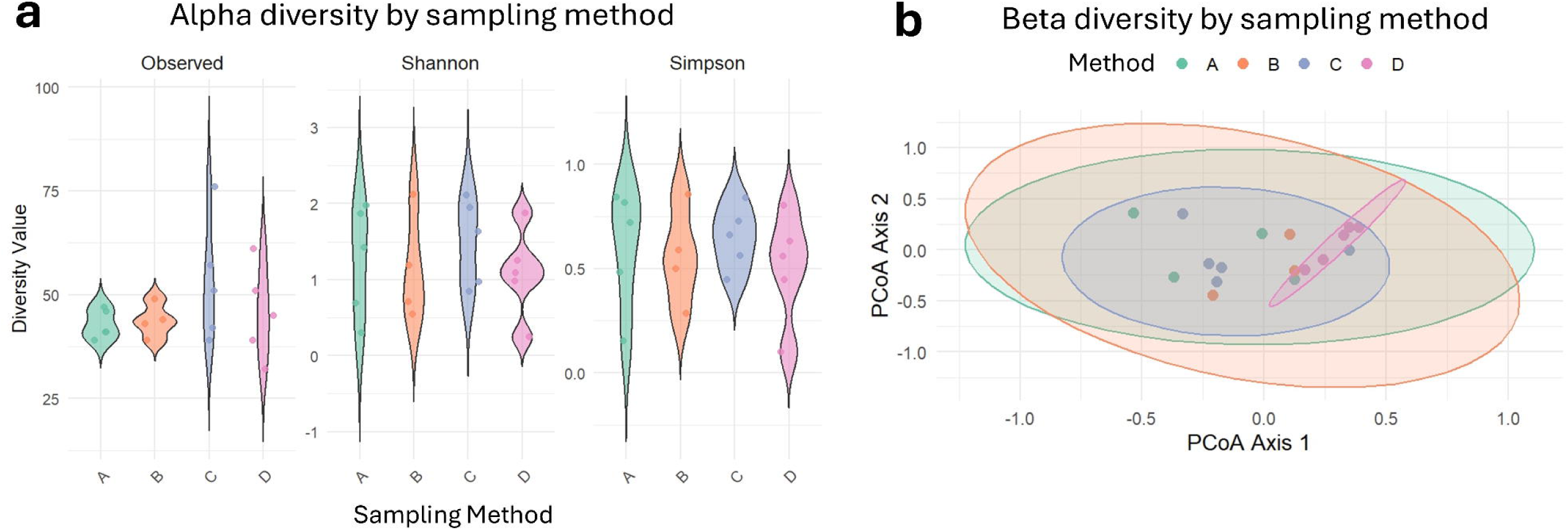
Microbial diversity across processing methods. (a) Violin plots showing alpha diversity metrics - Observed, Shannon, and Simpson indices, based on genus-level data across four processing methods (A-D). Each dot represents an individual sample, with colours corresponding to: Method A: green; Method B: orange; Method C: blue; Method D: pink. Samples were rarefied to 130,000 reads, and unassigned reads were excluded. A Kruskal-Wallis test revealed no statistically significant differences in alpha diversity between processing methods. (b) Principal Coordinate Analysis (PCoA) plots based on Bray-Curtis dissimilarity, comparing microbial community composition across the same four processing methods. Each dot represents an individual sample, and ellipses represent the 95% confidence interval for each method group. Samples were rarefied as above, and unassigned reads were excluded. Pairwise PERMANOVA tests indicated a significant difference between Method C and Method D (*p* = 0.028).

Beta diversity analysis using Bray-Curtis dissimilarity showed substantial overlap across all four methods (Figure 7b). Nevertheless, Methods A and B exhibited greater variability between samples, while Method D produced more consistent community profiles across patients. A pairwise PERMANOVA test revealed a statistically significant difference between Methods C and D, with processing method accounting for 22% of the observed variation (p = 0.028). Despite these differences, inter-patient variability was the dominant source of diversity, indicating that the choice of processing method had a minimal impact on overall microbial community structure (Figure S10).

### 3.7 Taxonomic composition by processing method

To assess the impact of sample processing on microbial community composition, taxonomic profiles were compared across the four methods (A-D) for five patients (Figure 8, S11). While overall diversity metrics showed minimal differences between methods, taxonomic assignment revealed method-specific variation in the dominant genera detected.

**Figure 8.**
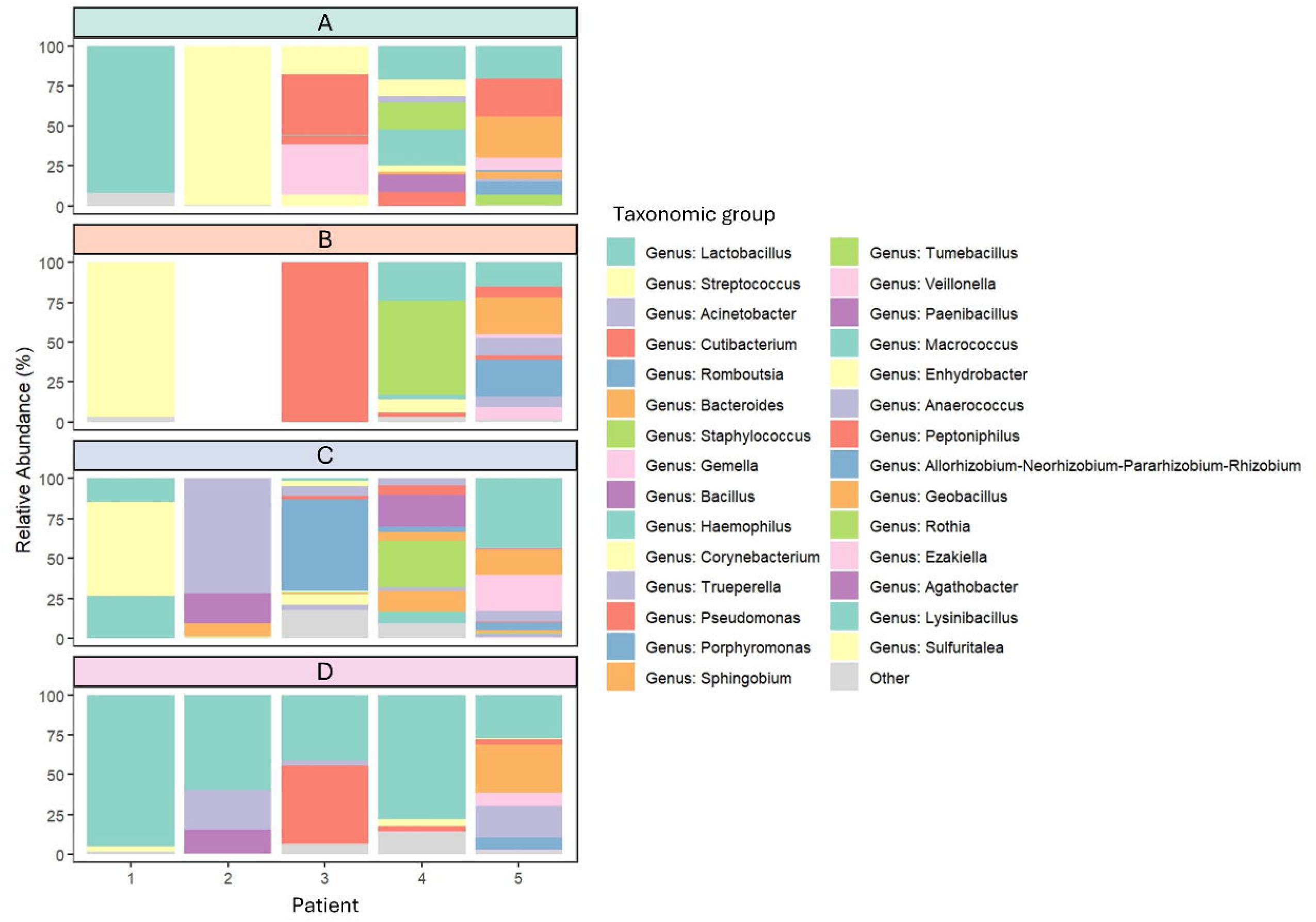
Relative abundance of taxonomic groups by patient and processing method. Bar plots showing the relative abundance of the 29 most abundant taxonomic groups within each sample. Taxonomic classification is presented at the genus level, or at a higher level when genus-level assignment was not possible. Unassigned reads were excluded from the analysis, and less abundant taxa were grouped under the category labelled “Other”. Samples represent five patients (1-5), each processed using four different methods (A-D). The sample from patient 2, processing method B, was excluded due to low read count.

In Patient 1, Methods A and D consistently identified *Lactobacillus* as the dominant genus, whereas Methods B and C primarily detected *Streptococcus*. In Patient 2, Method A identified *Corynebacterium* and *Streptococcus,* while Methods C and D detected *Acinetobacter*, with Method D also identifying *Lactobacillus*. Patient 3 showed further variation: Methods A, B, and D identified high levels of *Cutibacterium*, whereas Method C was dominated by *Romboutsia*. In contrast, Patient 5 demonstrated strong consistency across all four methods, with each identifying the same dominant genera, noting that this patient overall had a higher bacterial proportion in all swabs.

These findings suggest that while inter-patient variability is the primary driver of microbial composition, processing method can influence taxonomic resolution and detection sensitivity, particularly for low-abundance or method-sensitive taxa. Notably, Method D consistently identified *Lactobacillus* across multiple patients, a pattern not observed with other methods, indicating that freezing prior to processing may enhance recovery of certain genera.

Overall, minimal differences were observed between the four methods in terms of dominant taxa. However, Method D demonstrated a slightly higher bacterial read count and greater consistency in detecting key genera such as *Lactobacillus*. Given the sensitivity of low-biomass microbiome analyses to collection and processing variability, standardisation of methodology is essential. Taken together, Method D, which incorporates freezing prior to processing, was deemed to be optimal for low-biomass uterine swab analysis. It consistently yielded higher bacterial DNA concentrations, produced more stable community profiles, and maintained taxonomic resolution, making it well-suited for future studies of the uterine microbiome.

## 4. Discussion

This study provides foundational insights into the uterine microbiome of patients with endometrial cancer in New Zealand, addressing a significant gap in global microbiome research. Australasia remains markedly underrepresented in human microbiome studies, with most data originating from North America, Europe, and parts of Asia. Given that microbiome composition is influenced by geography, lifestyle, and ethnicity,^41^ findings from international cohorts cannot be directly extrapolated to New Zealand. This underscores the importance of region-specific investigations to uncover potentially unique microbial signatures relevant to disease pathogenesis and biomarker discovery.

### 4.1 Sequencing platform performance

Our comparison of Oxford Nanopore Technologies (ONT) and Illumina sequencing platforms revealed notable differences in microbial profiles, driven largely by the distinct 16S rRNA regions targeted by each platform. Prior studies have shown that different regions of the 16S gene preferentially detect specific taxa.^19, 42^ For example, the V1-V2 regions offer high sensitivity and specificity, while the V3-V4 regions better capture overall diversity. Combinations including the V3 region have demonstrated improved taxonomic resolution.^19, 43^

In our study, ONT sequencing of the full-length 16S rRNA gene (V1-V9) revealed greater richness and enabled more precise taxonomic assignments at lower levels (e.g., genus) compared to Illumina, which targeted only the V3 region. This limited region coverage likely contributed to Illumina’s reduced ability to assign reads at finer taxonomic levels.^44^ This enhanced resolution is particularly valuable in cancer research, where detecting low-abundance or method-sensitive taxa may reveal rare microbial signatures linked to tumour subtype, immune infiltration, or treatment response.

Additionally, the sequencing and bioinformatic pipelines differ significantly between platforms. ONT produced a higher read count, and alpha rarefaction analysis indicated that microbial diversity was not fully captured. In contrast, Illumina curves plateaued before 50,000 reads, suggesting saturation and lower overall richness. Although rarefying reads to equal depths is common practice, doing so here would have compromised the richness captured by ONT. Previous studies have reported mixed findings regarding platform performance, with some favouring Illumina and others ONT.^45, 46, 47^

Taxonomic assignment criteria also varied. Illumina assigned nearly all reads at the order level, while ONT achieved only ∼80% assignment at the phylum level. These differences stem from platform-specific error profiles and assignment algorithms.

Illumina uses a naïve Bayes classifier trained on the full 16S gene with a 99% similarity threshold, whereas ONT employs minimap2 with 95% similarity and 90% coverage thresholds, parameters chosen to accommodate ONT’s higher error rate.^15^ Despite these adjustments, ONT’s lower assignment rates at higher taxonomic levels likely reflect residual error and limited correction capabilities.^48^ However, ongoing improvements in ONT technology and bioinformatics pipelines continue to enhance accuracy.^15, 48^

Overall, ONT provided deeper resolution at lower taxonomic levels and a more comprehensive view of microbial communities in our uterine samples, consistent with other studies in low-biomass environments.^45, 46, 47, 49^

### 4.2 Species-level limitations and taxonomic ambiguity

Species-level identification using 16S rRNA sequencing remains challenging, particularly when using the SILVA database, which lacks robust species-level curation.^26, 50^ Accordingly, both Epi2me and QIIME2 advise against using SILVA for species-level assignments. Moreover, the 16S gene alone cannot reliably distinguish closely related species, such as those within *Shigella* and *Escherichia*.^51^ This limitation affects our ability to identify both clinically relevant species and exclude potential contaminants.

For instance, genera such as *Streptococcus*, *Staphylococcus*, and *Cutibacterium*, previously reported in uterine microbiome studies,^52, 53, 54, 55^ include species commonly found on skin, in soil, or in water. Without species-level resolution, it is difficult to determine whether these represent true uterine residents or contaminants. Similarly, *Porphyromonas somerae* has been implicated in endometrial cancer pathogenesis due to its potential role in promoting inflammation and epithelial disruption,^8, 56^ but while *Porphyromonas* was detected in most samples, we cannot confirm the presence of *P. somerae* specifically, limiting our ability to explore its association with tumour biology in our cohort without application of additional targeted methodology.

To overcome these limitations, future studies could incorporate whole metagenome shotgun sequencing or metatranscriptomics, which allow for species-level identification and functional profiling, but are more expensive and require more complex analysis pipelines than 16S rDNA analysis. These approaches could enable more precise characterisation of microbial communities and their metabolic or immunological contributions to EC. However, they require substantial sequencing depth and rigorous contamination control, particularly critical in low-biomass environments like the uterus. Statistical decontamination tools such as Decontam may also help distinguish true biological signals from artefactual noise, improving confidence in species-level associations with cancer outcomes.^38^

### 4.3 Uterine microbiome composition

The composition of the uterine microbiome in patients with and without endometrial cancer remains a subject of active investigation and debate, with evidence of global differences that are driven by methodological and cohort-specific factors, or a combination of both. While early studies suggested a dominance of *Lactobacillus* species in the healthy uterus, more recent research has revealed a more diverse and context-dependent microbial landscape, particularly in malignancy.

In our study, *Lactobacillus* was dominant in some samples, consistent with findings from,^11, 57, 58^ who reported *Lactobacillus crispatus* depletion in EC patients and its anti-proliferative effects on endometrial organoids. However, other samples showed enrichment of taxa such as *Acinetobacter*, *Pseudomonas*, and *Comamonadaceae*, aligning with Winters, Romero ^59^ findings in EC patients. These genera are often associated with environmental niches and may reflect either true colonisation or contamination, warranting expansion of global uterine datasets to answer this ongoing question.

Beyond these genera, our dataset also detected *Bacteroides*, *Prevotella,* and *Shigella*, which have been previously linked to EC and other gynaecological conditions.^9, 10^ *Prevotella* and *Porphyromonas*, in particular, have emerged as recurrent taxa in EC-associated microbiomes. *Porphyromonas somerae*, for example, has been shown to invade endometrial cells and exhibit pathogenic behaviour stimulated by oestrogen exposure, a key risk factor in EC.

Other studies have identified *Anaerococcus*, *Peptoniphilus*, and *Buttiaxella* as enriched in EC patients, with microbial swapping between vaginal and rectal sites potentially contributing to uterine colonisation.^60^ These taxa may influence inflammation, immune modulation, and hormone metabolism, processes central to EC pathogenesis. Moreover, the vaginal microbiome has been shown to correlate with tumour grade and histology, with distinct community state types predictive of low-grade versus high-grade disease.^61^

Taken together, our findings are broadly consistent with previous reports, but also highlight the complexity and variability of the uterine microbiome in EC. The presence of both protective (*Lactobacillus*) and potentially pathogenic (*Porphyromonas*, *Prevotella*) taxa suggests that microbial balance may influence disease trajectory. However, distinguishing true residents from contaminants remains a challenge, particularly in low-biomass environments.

To fully characterise the endometrial cancer microbiome in New Zealand and beyond, future studies will require larger, ethnically diverse cohorts, longitudinal sampling, and integration of functional data (e.g., metagenomics, metabolomics). Such efforts could uncover microbial biomarkers for early detection, stratify patients by risk, and inform microbiome-targeted interventions in EC care.

### 4.4 Contamination risks and control strategies

Contamination remains one of the most significant challenges in low-biomass microbiome studies, where even small amounts of exogenous bacterial DNA can dominate sequencing results, masking true microbial signals and potentially leading to false associations with disease.^12^ Erroneous findings can also arise during analysis by mis-annotation of bacterial sequences.^62^ This is particularly problematic in cancer research, where microbial signatures are increasingly being explored as diagnostic or prognostic biomarkers. Misidentifying contaminants as disease-associated taxa could mislead biomarker discovery efforts and compromise translational applications, underscoring the importance of rigorous contamination control throughout the experimental workflow.^63, 64^

Contamination sources are diverse and include sampling instruments, DNA extraction kits, sequencing reagents, and laboratory environments. Commercial DNA extraction kits carry their own “kit microbiome”, which can introduce consistent but misleading microbial profiles.^12, 65^ In the context of endometrial cancer, where microbial loads are low and the clinical implications of microbial shifts are still being defined, even minor contamination can have outsized effects on interpretation. Therefore, a robust contamination testing strategy must incorporate collection-site swabs, DNA extraction blanks, and no-template sequencing controls, each tailored to the specific reagents and protocols used at every stage.^39^

Our study incorporated negative controls, including DNA extraction blanks and no-template sequencing controls, to monitor potential contamination at key stages of the ONT workflow. Taxa typically associated with human skin (*Cutibacterium*, *Enhydrobacter*), water (*Acinetobacter*), and vaginal flora (*Lactobacillus*) were occasionally detected in these controls. These observations informed our criteria for interpreting microbial profiles, helping to distinguish genuine signals from possible background contributions.^63^ To improve confidence in microbial signals and support robust biomarker discovery, future studies would benefit from broader and stage-specific controls, alongside computational decontamination tools and refined protocols to minimise background interference.

### 4.5 Sample processing methods and reproducibility

Variations in sample collection and storage methods contribute to subtle differences in microbial profiles. These could stem from differences in contamination risk during collection and processing, including the use of distinct swab types, storage conditions, laboratory equipment, and environmental exposure.^12^ Additionally, methodological differences, such as using the cell pellet (samples C and D) versus the whole sample (A and B), and the impact of freezing, clearly influenced microbial composition in our study.

Freezing can disrupt bacterial cell walls, particularly in Gram-positive species, enhancing DNA extraction efficiency. In faecal microbiome studies, this has been associated with an increased *Firmicutes*-to-*Bacteroidetes* ratio.^66, 67^ However, no such trend was observed in our data. While the differences between the four collection methods were not unexpected, they highlight the importance of consistency in DNA collection and storage protocols when comparing across experiments or studies.^68^

With relevance to endometrial cancer, reproducibility in microbiome profiling is essential for identifying microbial biomarkers that could inform diagnosis, prognosis, or therapeutic response. Methodological inconsistencies can obscure biologically meaningful patterns and hinder cross-study comparisons. Standardising protocols for low-biomass sample handling will be critical for advancing microbiome research in EC and enabling robust meta-analyses across diverse cohorts.

This pilot study lays the groundwork for future investigations into the uterine microbiome in endometrial cancer, particularly within underrepresented populations such as those in New Zealand. While we did not aim to fully characterise the microbial landscape associated with endometrial cancer, we successfully established a reproducible protocol for uterine sample collection and processing that improves bacterial DNA recovery and sequencing depth. Our comparative analysis identified ONT as the preferred platform for low-biomass environments, offering superior genus-level resolution and broader taxonomic coverage. Importantly, we demonstrated the necessity of implementing contamination controls throughout the workflow, from sample collection to sequencing, to ensure accurate microbial profiling. This is especially critical in cancer research, where microbial signals may be subtle yet biologically significant.

Ultimately, a clearer understanding of the uterine microbiome may inform diagnostic or therapeutic strategies for endometrial cancer. Microbial signatures could serve as biomarkers for early detection, stratify patients by tumour subtype or risk, or even guide microbiome-targeted interventions. Future research should explore longitudinal sampling, integrate host transcriptomic or immunologic data, and assess microbial function to better understand host–microbiome interactions in endometrial cancer.

## Ethical Statement

Ethics approval was obtained from the New Zealand Health and Disability Ethics Committee (HDEC 15/CEN143 and HDEC 2022 EXP 12616) and the University of Otago (H20/002 and H25/0421).

## Data Availability Statement

The data that support the findings of this study are openly available in the NCBI Sequence Read Archive (SRA) at https://www.ncbi.nlm.nih.gov/bioproject/PRJNA1313862, reference number PRJNA1313862, and will be available following an embargo from the date of publication. The data are scheduled for public release on 27 September 2026.

## Acknowledgements

This work was supported by the Cancer Research Trust New Zealand, the Cancer Society New Zealand, and the Maurice and Phyllis Paykel Trust. We are deeply grateful to the women who generously participated in this biobanking study and made this research possible.

The authors also acknowledge the use of the New Zealand eScience Infrastructure (NeSI) high-performance computing facilities, consulting support, and training services as part of this research. New Zealand’s national facilities are provided by NeSI and funded jointly by NeSI’s collaborator institutions and through the Ministry of Business, Innovation & Employment’s Research Infrastructure programme (https://www.nesi.org.nz). We also thank the team at Auckland Genomics, University of Auckland, for their expertise and assistance with PCR and sequencing.

## Authors’ Contributions

*Conceptualisation*, C.B., A.A., C.H.; *Methodology*, C.B., A.A.; *Investigation*, A.A., S.B., B.N.; *Sample and Clinical Data Collection*, C.H.; *Software*, S.B.; *Formal Analysis*, S.B.; *Resources*, C.H., S.F., A.A., C.B.; *Data Curation,* S.B.; *Writing - Original Draft*, S.B., A.A., C.B.; *Visualisation*, S.B; *Supervision*, A.A., C.B; *Funding Acquisition*, A.A., C.B.

