## Supplemental Figure 1 for "Evaluating sequencing strategies for endometrial microbiome profiling in endometrial cancer: a comparative study of short- and long-read 16S rRNA approaches"

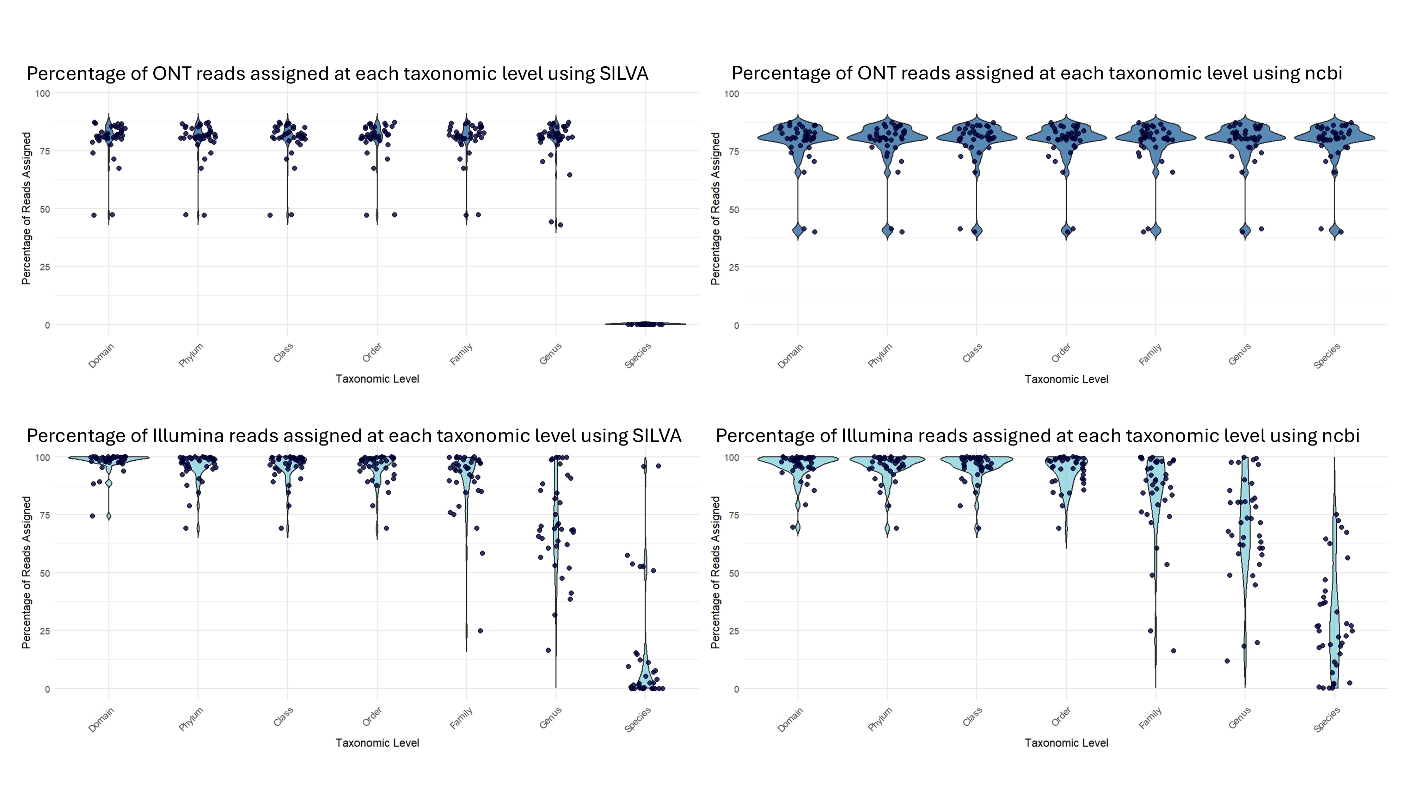

**Figure S1.** Taxonomic assignment of sequencing reads across platforms and databases. Violin plots showing the percentage of reads assigned at each taxonomic level for samples sequenced using ONT and Illumina platforms. Taxonomic classification was performed using both the NCBI and SILVA databases. Each dot represents an individual sample (n = 38). Reads classified as unknown, unassigned, eukaryotic, or uncultured were excluded from the assigned read count.
