## Supplemental Figure 2 for "Evaluating sequencing strategies for endometrial microbiome profiling in endometrial cancer: a comparative study of short- and long-read 16S rRNA approaches"

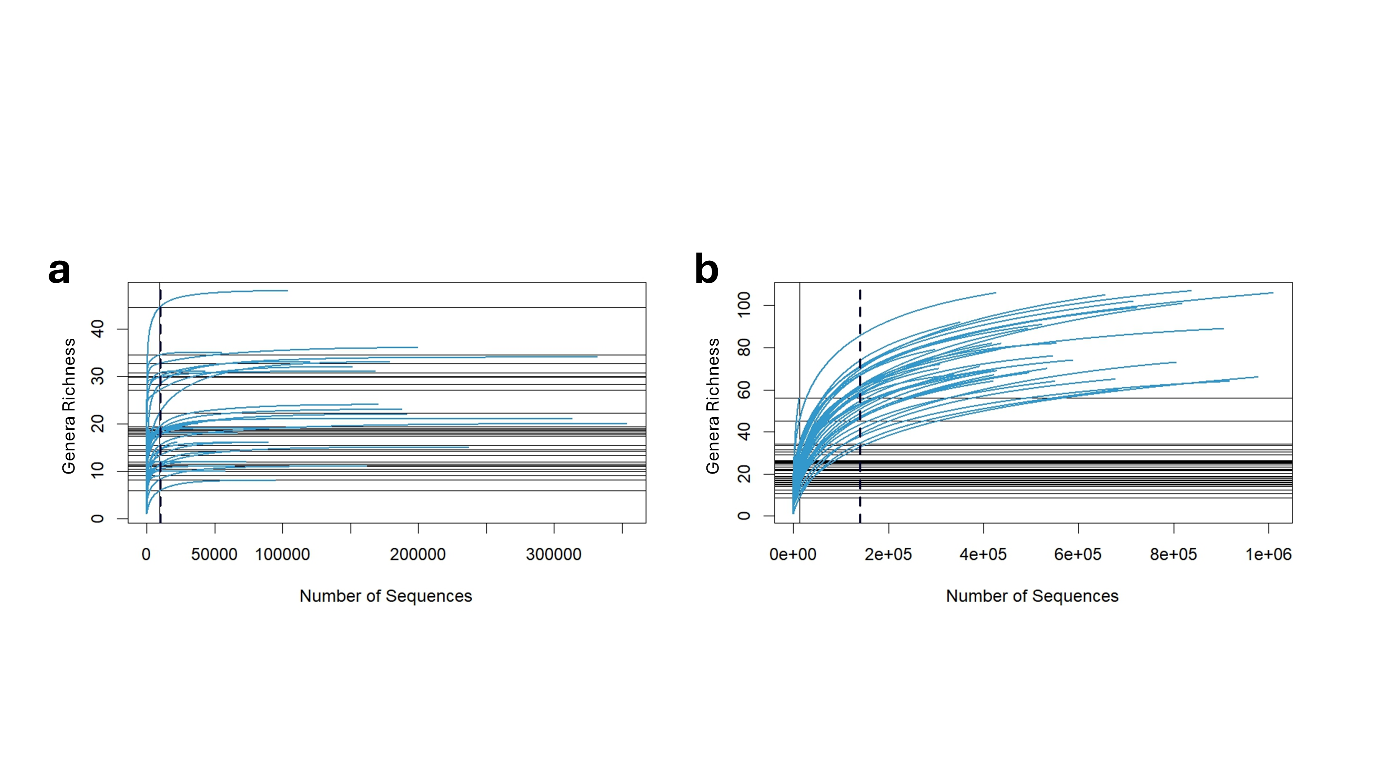

**Figure S2.** Alpha rarefaction analysis of Illumina and ONT sequencing data. Alpha rarefaction plots showing genus-level microbial diversity for (a) Illumina and (b) Oxford Nanopore Technologies datasets. The dotted line indicates the read depth used for diversity analysis: 10,000 reads for Illumina and 140,000 reads for ONT.
