## Supplemental Figure 3 for "Evaluating sequencing strategies for endometrial microbiome profiling in endometrial cancer: a comparative study of short- and long-read 16S rRNA approaches"

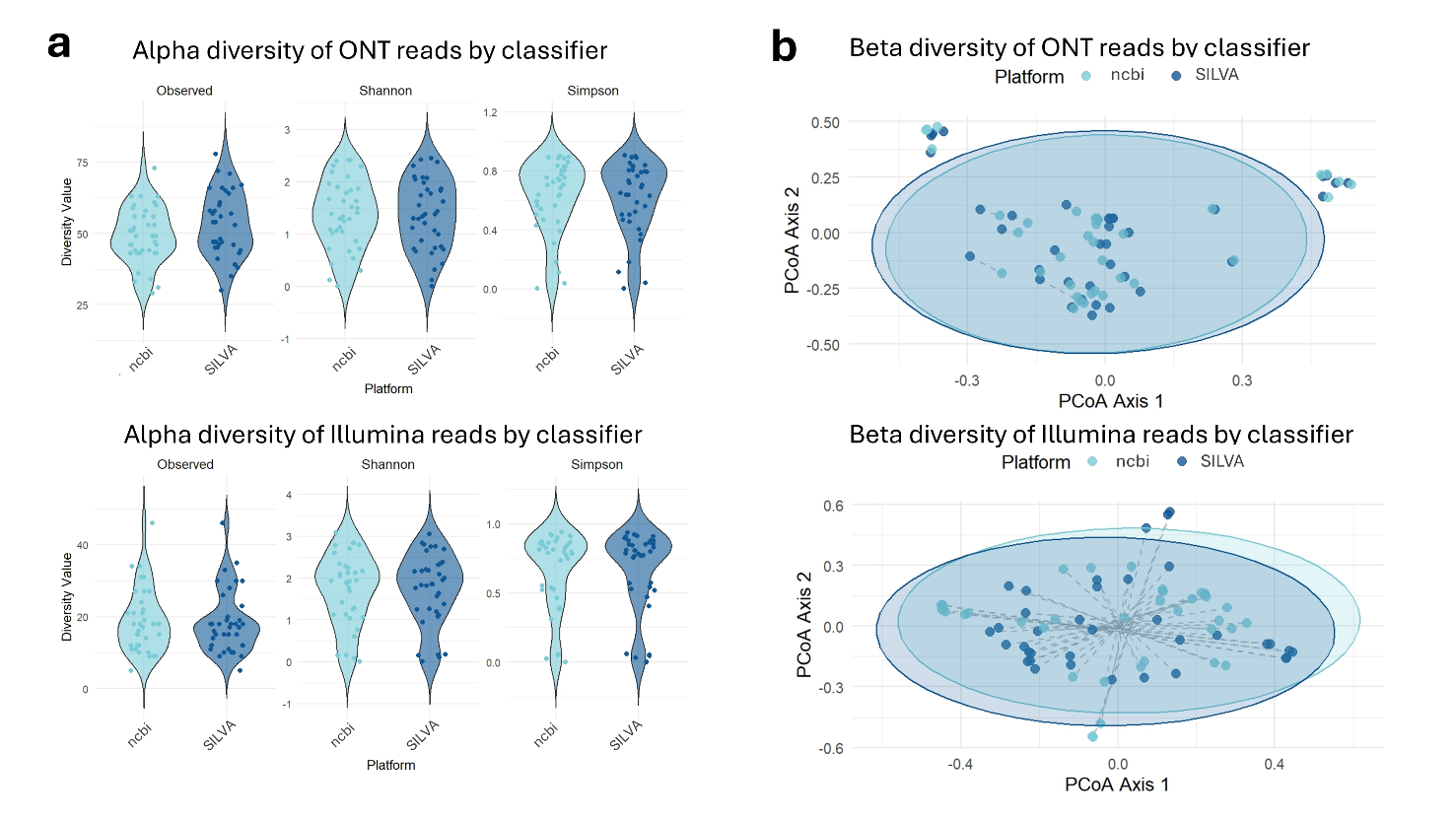

**Figure S3.** Comparison of alpha and beta diversity metrics by taxonomic database. (a) Violin plots showing alpha diversity metrics - Observed, Shannon, and Simpson indices, based on genus-level data. Taxonomy was assigned using either the NCBI (light blue) or SILVA (dark blue) databases. Each dot represents an individual sample. Samples were rarefied to 10,000 reads for Illumina and 140,000 reads for ONT; unassigned reads were excluded. Wilcoxon tests revealed no statistically significant differences in diversity metrics between databases within the same sequencing platform. (b) Principal Coordinate Analysis (PCoA) plots based on Bray-Curtis dissimilarity, comparing genus-level taxonomic assignments using NCBI (light blue) and SILVA (dark blue). Each dot represents an individual sample, with dotted lines connecting the same sample classified by both databases. Ellipses represent the 95% confidence interval for each database group. Samples were rarefied as above, and unassigned reads were excluded. PERMANOVA tests indicated significant differences in community composition between databases: ONT: *F* = 8.19, *p* = 0.001 (999 permutations); Illumina: *F* = 11.2, *p* = 0.001.
