## Supplemental Figure 4 for "Evaluating sequencing strategies for endometrial microbiome profiling in endometrial cancer: a comparative study of short- and long-read 16S rRNA approaches"

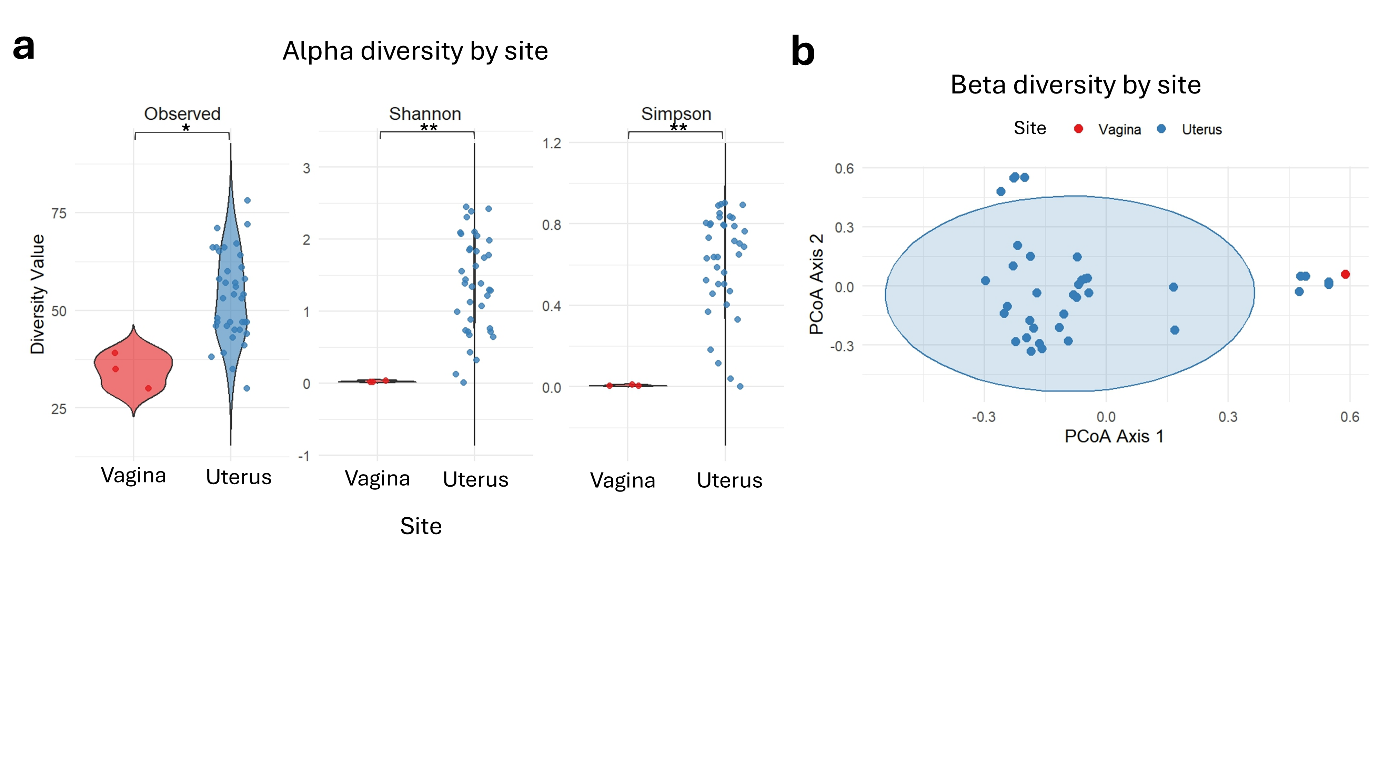

**Figure S4.** Comparison of microbial diversity by sample site. (a) Violin plots showing alpha diversity metrics - Observed, Shannon, and Simpson indices, based on genus-level data, comparing samples from the uterus (blue, *n* = 38) and vagina (red, *n* = 3). Each dot represents an individual sample. Samples were rarefied to 140,000 reads, and unassigned reads were excluded. Statistical significance was assessed using the Wilcoxon test: *p* < 0.05 (*), *p* < 0.01 (**). (b) Principal Coordinate Analysis (PCoA) plots based on Bray-Curtis dissimilarity, comparing microbial communities between uterus (blue, *n* = 38) and vagina (red, *n* = 3) swabs at the genus level. Each dot represents an individual sample. The ellipse indicates the 95% confidence interval for uterus samples. Samples were rarefied as above, and unassigned reads were excluded. PERMANOVA test results: *F* = 3.08, *p* = 0.003 (999 permutations).
