## Supplemental Figure 5 for "Evaluating sequencing strategies for endometrial microbiome profiling in endometrial cancer: a comparative study of short- and long-read 16S rRNA approaches"

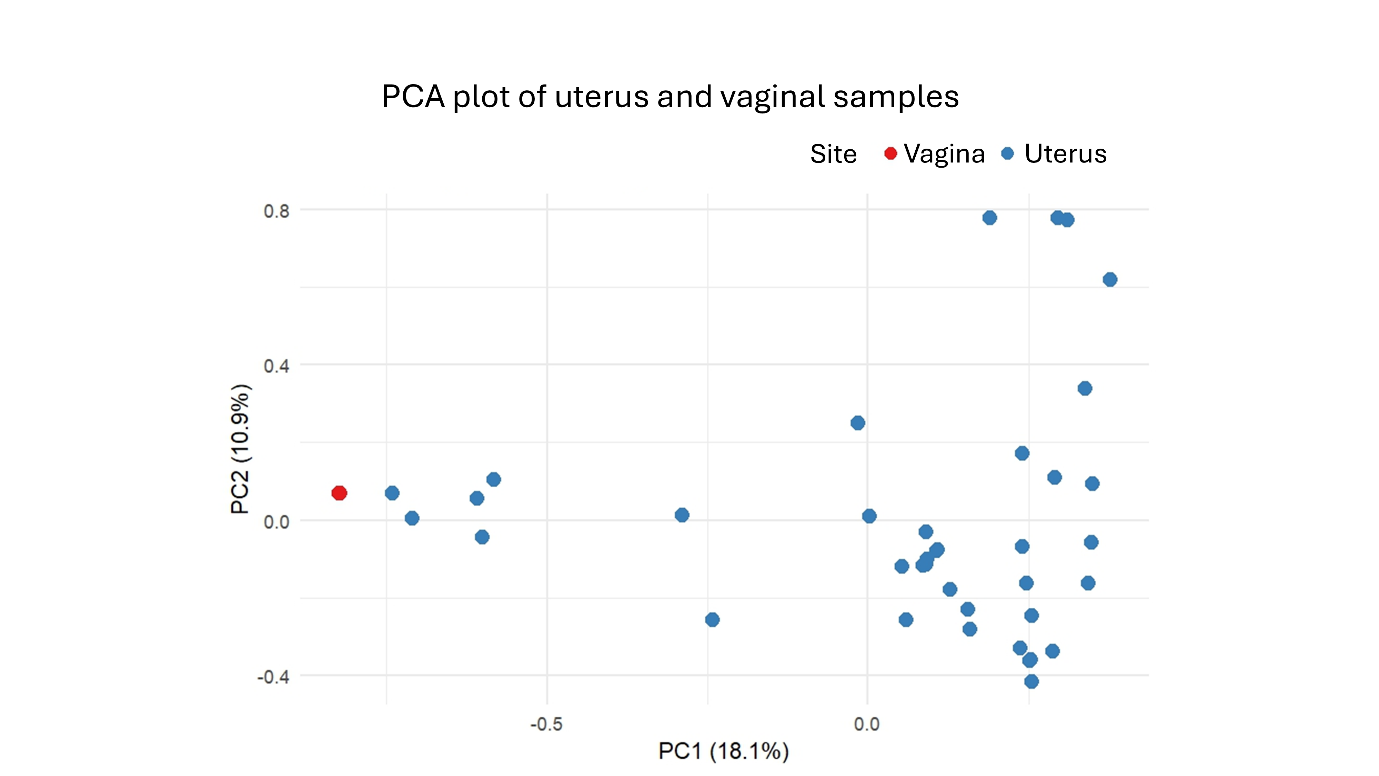

**Figure S5.** Principal component analysis of ONT samples by sample site. Principal Component Analysis (PCA) plot based on Hellinger-transformed genus-level data from ONT sequencing. Samples were derived from either the uterus (blue, *n* = 38) or vagina (red, *n* = 3).
