## Supplemental Figure 6 for "Evaluating sequencing strategies for endometrial microbiome profiling in endometrial cancer: a comparative study of short- and long-read 16S rRNA approaches"

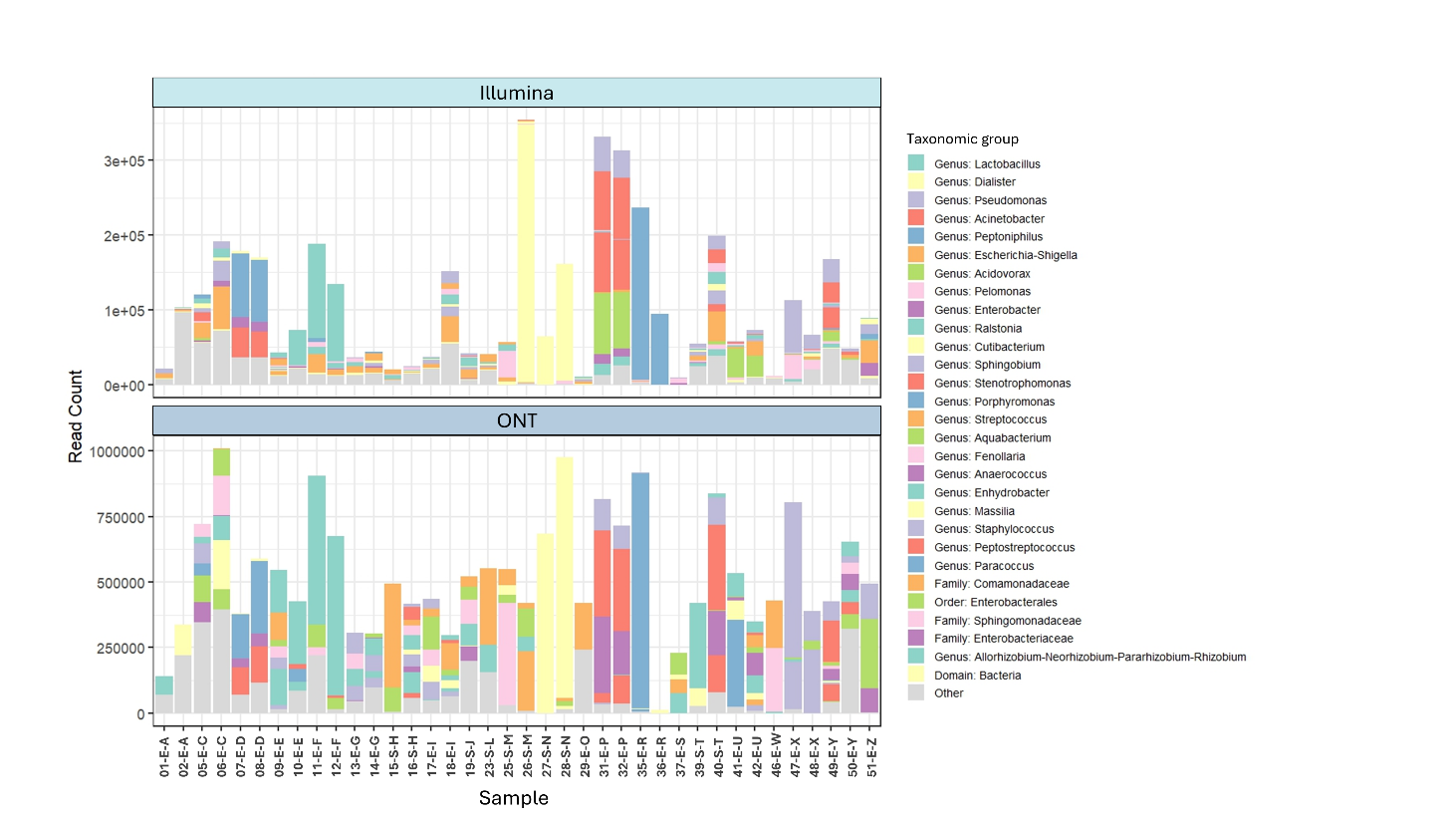

**Figure S6.** Taxonomic composition of samples based on read abundance. Bar plots showing the read counts of the 29 most abundant taxonomic groups within each sample. Taxonomic classification was performed at the genus level, or at a higher taxonomic level when genus-level assignment was not possible. Taxa with lower abundance were grouped into a single category labelled “Other” to simplify visualisation.
