## Supplemental Figure 7 for "Evaluating sequencing strategies for endometrial microbiome profiling in endometrial cancer: a comparative study of short- and long-read 16S rRNA approaches"

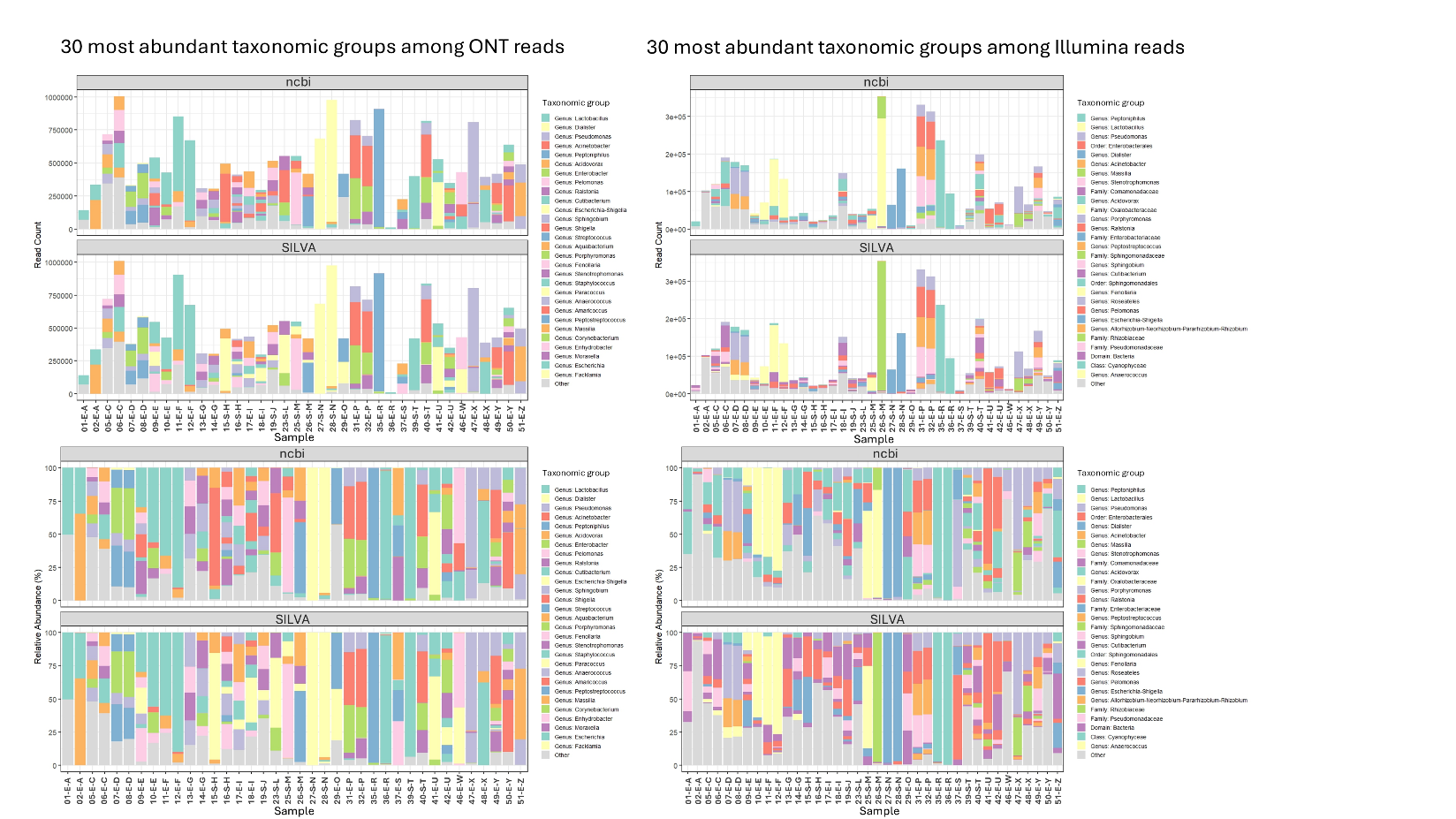

**Figure S7.** Comparison of taxonomic composition across sequencing platforms and databases. Bar plots showing the read counts of the 29 most abundant taxonomic groups within each sample. Samples sequenced using ONT are displayed on the left, and those sequenced using Illumina on the right. Taxonomic classification was performed using both the SILVA and NCBI databases. For each sample, the read count (top panels) and relative abundance (bottom panels) of taxonomic groups are shown. Taxa with lower abundance were grouped into a single category labelled “Other”. Taxonomic groups are presented at the genus level, or at a higher level when genus-level assignment was not possible. Unassigned reads were excluded from the analysis.
