## Supplemental Figure 8 for "Evaluating sequencing strategies for endometrial microbiome profiling in endometrial cancer: a comparative study of short- and long-read 16S rRNA approaches"

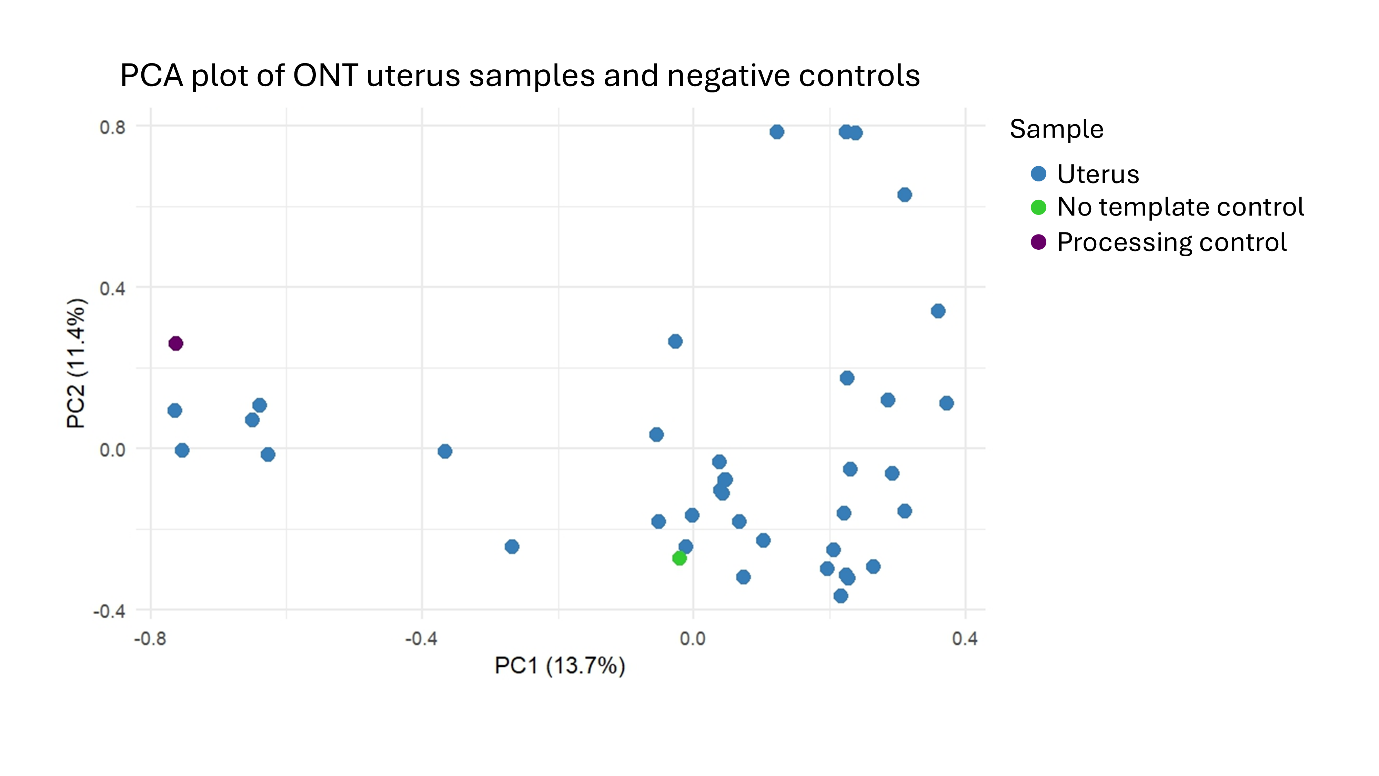

**Figure S8.** Hellinger-transformed PCA of uterus samples and negative controls. Principal Component Analysis (PCA) plot based on Hellinger-transformed genus-level data from ONT sequencing. The plot compares uterus samples collected for platform comparison (blue, *n* = 38) with two types of negative controls: no template control (green) added during the sequencing process and processing control (purple) added prior to DNA isolation and quantification.
