## Supplemental Figure 9 for "Evaluating sequencing strategies for endometrial microbiome profiling in endometrial cancer: a comparative study of short- and long-read 16S rRNA approaches"

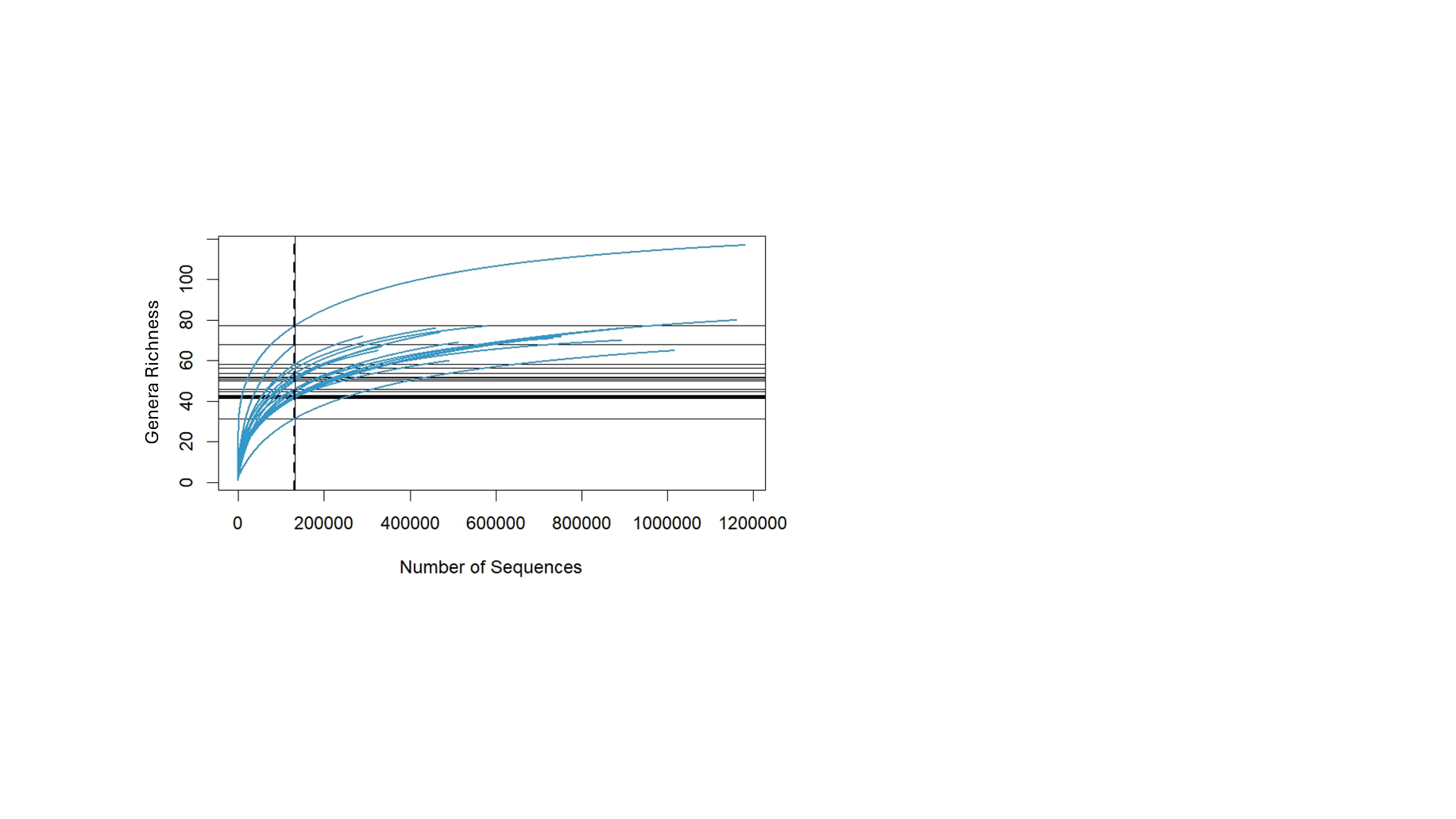

**Figure S9.** Alpha rarefaction analysis of ONT samples by collection and storage method. Alpha rarefaction plots showing genus-level microbial diversity in ONT-sequenced samples used to compare different collection and storage methods. The dotted line at 130,000 reads indicates the read depth used for diversity analysis.
