## Supplemental Figure 10 for "Evaluating sequencing strategies for endometrial microbiome profiling in endometrial cancer: a comparative study of short- and long-read 16S rRNA approaches"

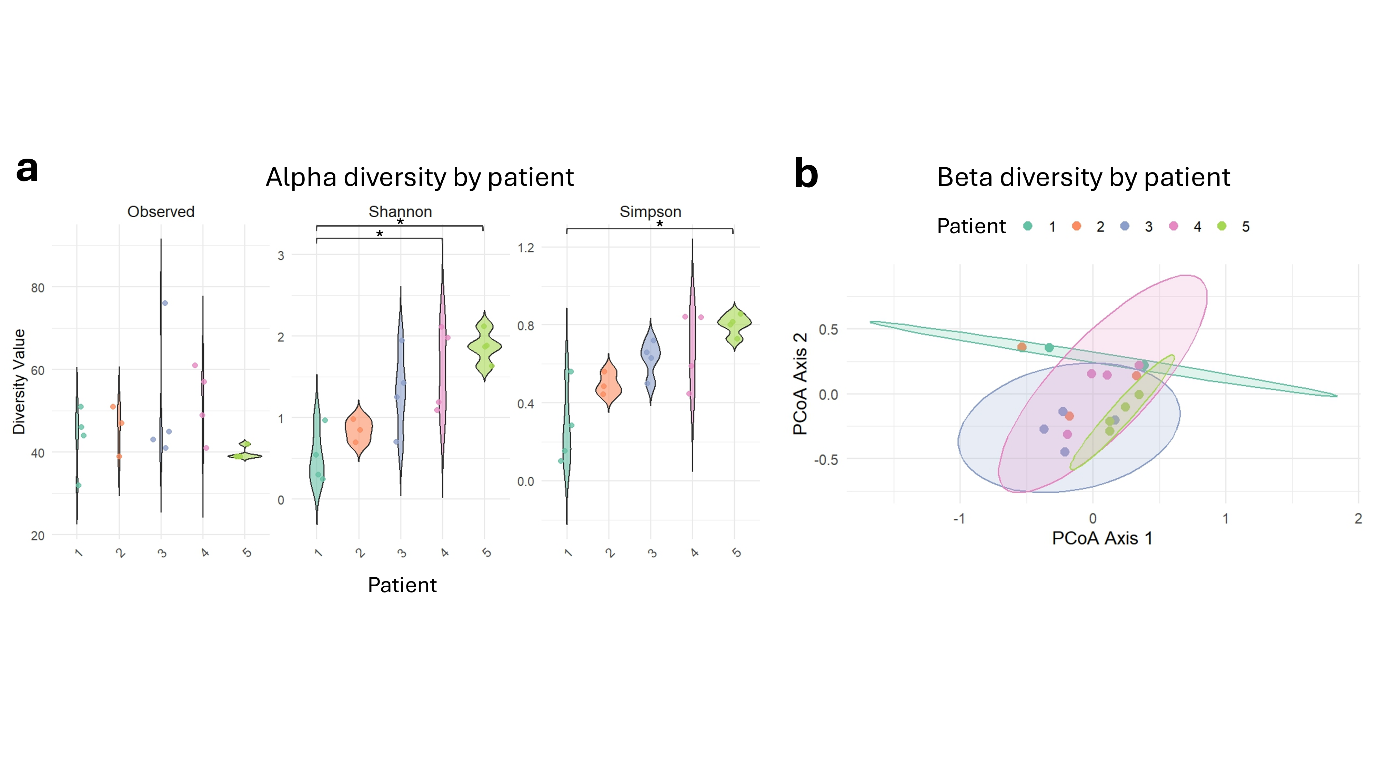

**Figure S10.** Microbial diversity across patients using ONT sequencing. (a) Violin plots showing alpha diversity metrics - Observed, Shannon, and Simpson indices, based on genus-level data for samples collected from patients 1-5. Each dot represents an individual sample. Samples were rarefied to 130,000 reads, and unassigned reads were excluded. Statistical significance was assessed using the Kruskal-Wallis test, followed by Dunn’s post hoc test. *p* < 0.05 (*). (b) Principal Coordinate Analysis (PCoA) plots based on Bray-Curtis dissimilarity, comparing microbial community composition across patients 1-5 at the genus level. Each dot represents an individual sample. Ellipses represent the 95% confidence interval for each patient group, excluding patient 2, which had too few samples for ellipse calculation. Samples were rarefied as above, and unassigned reads were excluded. Pairwise PERMANOVA tests revealed significant differences between: Patients 1 and 5 (*p* = 0.029), Patients 2 and 5 (*p* = 0.035), Patients 3 and 5 (*p* = 0.028), Patients 4 and 5 (*p* = 0.028).
