## Supplemental Figure 11 for "Evaluating sequencing strategies for endometrial microbiome profiling in endometrial cancer: a comparative study of short- and long-read 16S rRNA approaches"

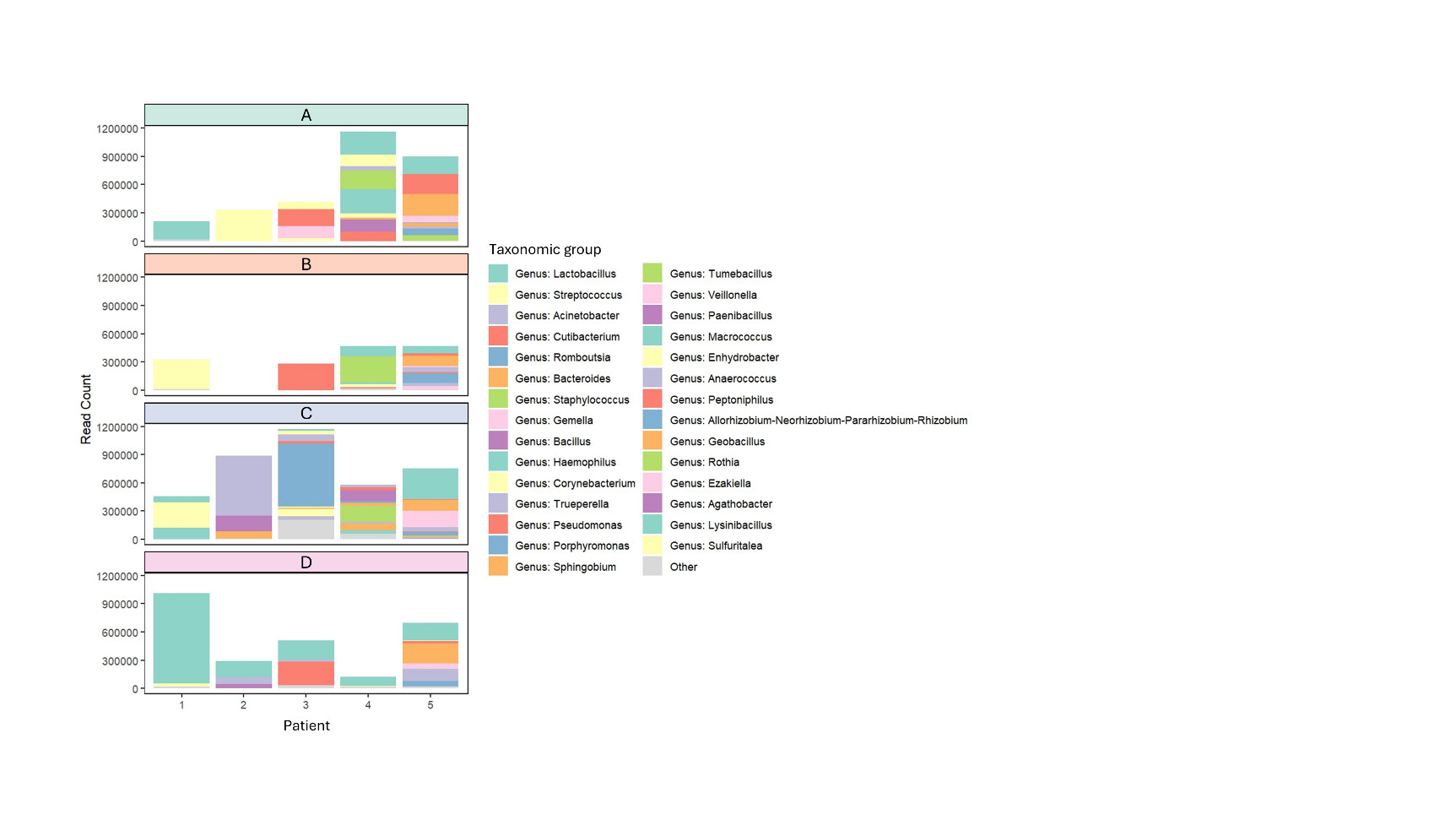

**Figure S11.** Taxonomic composition of samples by patient and processing method. Bar plots showing the read counts of the 29 most abundant taxonomic groups within each sample. Taxonomic classification is presented at the genus level, or at a higher level when genus-level assignment was not possible. Unassigned reads were excluded from the analysis, and less abundant taxa were grouped under the category labelled “Other”. Samples represent five patients (1-5), each processed using four different methods (A-D). The sample from patient 2, processing method B, was excluded due to low read count.
